# Type 2 innate immunity regulates hair follicle homeostasis to control *Demodex* pathosymbionts

**DOI:** 10.1101/2021.03.29.437438

**Authors:** Roberto R. Ricardo-Gonzalez, Maya E. Kotas, Iliana Tenvooren, Diana M. Marquez, Marlys S. Fassett, Jinwoo Lee, Scott G. Daniel, Kyle Bittinger, Roberto Efraín Díaz, James S. Fraser, K. Mark Ansel, Matthew H. Spitzer, Hong-Erh Liang, Richard M. Locksley

## Abstract

Allergic skin diseases are common, but basal roles for type 2 immunity in cutaneous homeostasis are incompletely understood. Here, we show that skin group 2 innate lymphoid cells (ILC2s) are the predominant resident cells that secretes IL-13, which attenuates epithelial cell proliferation during anagen throughout the hair follicle (HF), including in HF stem cells. Although HF are normal in the absence of type 2 immunity, colonization with the commensal mite, *Demodex musculi,* results in epithelial proliferation and aberrant hair follicle morphology accompanied by loss of stable commensalism with massive infestation and expansion of inflammatory ILC2s and other immune cells. Topical anti-parasitic agents, but not broad-spectrum antibiotics, reversed the phenotype. Commensal *Demodex* colonization of mammalian hair follicles is ubiquitous, including in humans, revealing an unanticipated role for ILC2s and type 2 immunity for skin homeostasis in the normal environment.

**One Sentence Summary:** Dynamic regulation of skin physiology by Type 2 innate immunity regulates healthy commensalism by parasitic *Demodex* mites.

## Main

Type 2 immunity underlies allergic pathology accompanying atopic dermatitis (AD), food allergy and asthma, but also promotes epithelial homeostasis and repair in response to inflammation, including by pathogens^1, 2^. The discovery of innate group 2 lymphoid cells (ILC2s), which reside and propagate from embedded precursors in tissues as resident immune cells^3^, creates the opportunity to probe the basal state of type 2 immunity and perhaps better understand the increasing prevalence of allergic diseases. Type 2 cytokine dysregulation plays a central role in prevalent human skin diseases like AD, as corroborated by the efficacy of targeting the IL-4/IL-13 pathway^4^. With this in mind, we examined whether activated ILC2s play a role in normal cutaneous biology, revealing an unanticipated contribution to sustaining commensalism of an ubiquitous colonizer of mammalian hair follicles (HF).

### ILC2s are dynamically activated in skin

To investigate the function of skin ILC2s, we used previously characterized reporter mice for the type 2 cytokines IL-5 and IL-13^5, 6^ to assess the localization and functional status of ILC2s in resting skin. Corroborating our previous findings^7, 8^, ILC2s were the most prevalent cells expressing these cytokines under basal conditions (Fig. 1a and Supplementary Fig. 1), and IL-5-expressing cells localized predominantly within the epithelial layer in close association with HF (Fig. 1b–d). The positioning of activated ILC2s suggested the possibility that these cells might influence the homeostatic oscillation of growth and rest occurring in HF^9^. We observed that ILC2s activate cyclically, as assessed by IL-13 expression, which is high during the early postnatal period (days 1-14; when ILC2s activate^8^) and then subsequently undergoes oscillating bursts of expression characterized by low IL-13 during telogen (rest) and high IL-13 during anagen (growth) phases of the hair cycle, which remains synchronous in the first months of life (Fig. 1e). In order to confirm our IL-13 reporter mice were an accurate reflection of *Il13* gene expression, we sorted ILC2s from back skin of mice in telogen or anagen phase of the hair cycle, and independently confirmed the increase in *Il13* during anagen by quantitative PCR (Fig. 1f). Taken together, these observations suggest that ILC2s and IL-13 may function to support the cyclic growth and turnover of the HF compartment.

**Fig. 1.**
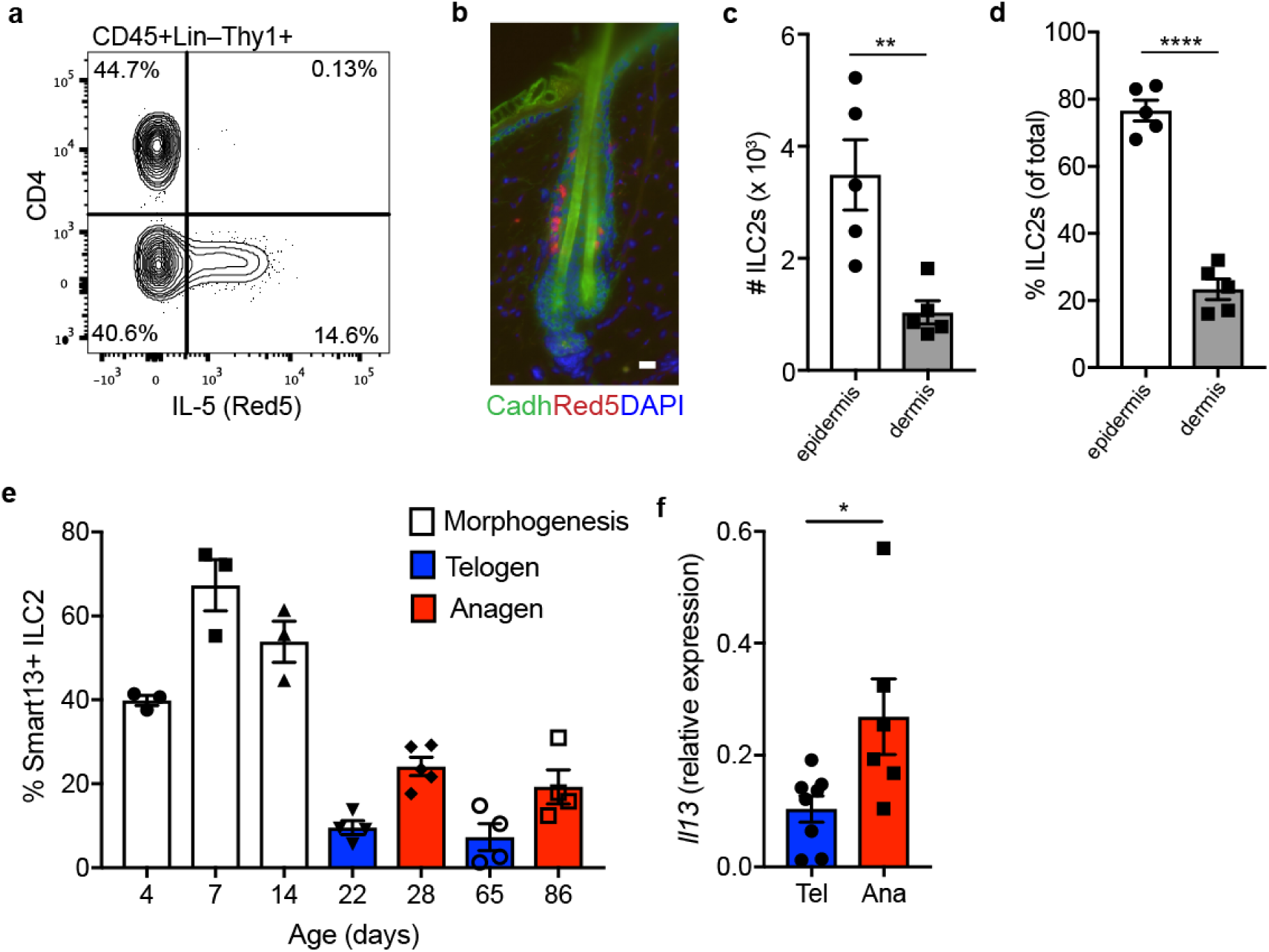
ILC2s in skin are a critical source of type 2 cytokines. **a**, Flow cytometry panel (pre-gated CD45^+^Lin^−^Thy1^+^) of IL-5 expression by CD4^+^ cells or ILC2s in adult mouse skin at homeostasis. **b**, Localization of IL-5 producing ILC2s (red) within the skin epithelium. E-cadherin (green) is used to highlight hair follicles. DAPI is represented in blue. Scale bar, 20 μm. **c**, Quantification of ILC2s (CD45^+^Lin^−^Thy1^+^Red5^+^) in the epidermis and dermis of adult mice. **d**, Percentage of ILC2s localized to the dermis or epidermis in *Il5^Red5/+^*. **e**, Quantification of expression of IL-13 by ILC2s throughout the various stages of the hair follicle cycle. **f**, Quantitative PCR for *Il13* expression by ILC2s (CD45+Lin–Thy1+Red5+) sorted from mice back skin in telogen or anagen stage of the hair cycle. Data presented as mean ± s.e.m. Data are representative of at least two independent experiments with n=3 or more mice per individual group/time point. *P < 0.05, **P < 0.01, ***P < 0.001, and ****P < 0.0001 as calculated with two-tailed Student’s t test.

### Type 2 cytokines impede proliferation throughout the epithelial compartment

To assess the role of IL-13 during anagen, we administered IL-13 or IL-4, which share the common IL-4Rα receptor, after depilation used to induce anagen and synchronous hair regrowth^10, 11^. Either IL-13 or IL-4, administered as long-lived antibody complexes, slowed hair regrowth (Fig. 2a,b), and was accompanied by a reduction in proliferation of HF stem cells (HFSCs) (Fig. 2c,d). We used mass cytometry (CyTOF) to assess the impact of IL-13/4 signaling on cutaneous epithelia. We observed that type 2 cytokines decrease the proliferation of the epithelial fraction as assessed by a decrease in Ki-67 and reduction in the uptake of the EdU analog, 5-Iodo-2-deoxyuridine (IdU) (Supplementary Fig. 2a). Conversely, when we characterized our IL-4Ra^−/−^ mice, which lack the capacity to respond to IL-4 and IL-13, we noted increased proliferation across the epithelial compartment at homeostasis (Supplementary Fig. 2a). Of note, epithelial populations responded broadly to type 2 cytokines as assessed by detection of phospho-STAT6, whereas disruption of the type 2 axis revealed aberrant activation of phospho-STAT1 and phospho-STAT4, and decreased phospho-STAT3 (Supplementary Fig. 2b), the latter previously implicated in maintaining skin wound healing and repair^12^.

**Fig. 2.**
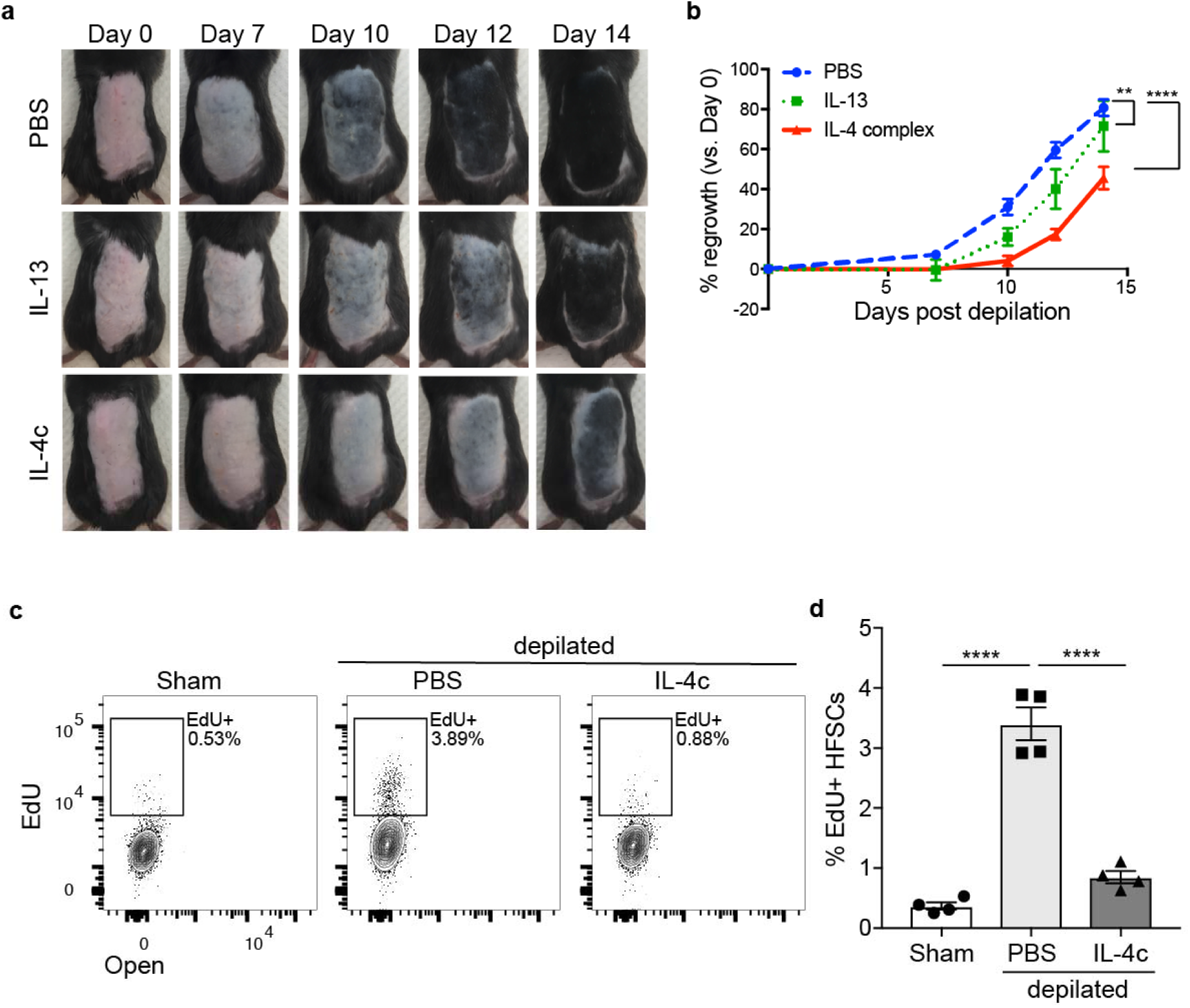
Type 2 cytokines slow hair regrowth and decrease HFSC proliferation. **a**, Effect of IL-4 and IL-13 subcutaneous injection in hair growth. Wild type C57BL/6 mice were treated with PBS, IL-13 or IL-4 complex (IL-4c) on the first 2 days after depilation (Days 0 and 1) and their hair regrowth pattern was tracked over 2 weeks. One representative mouse per treatment arm is shown. **b**, Quantification of hair regrowth from PBS, IL-4c, or IL-13 treated mice as in (A). **c**, Representative flow cytometry panel of hair follicle stem cells (HFSCs, pre-gated as CD45^−^MHCII^−^CD34^+^Alpha6^+^) after EdU. Mice were depilated as in (**a**) and treated with IL-4 complex (IL-4c) on the first two days after depilation. Mice were harvested on day 4 after depilation. **d**, Percent EdU^+^ HFSCs four days after depilation as in (**c**). Data presented as mean ± s.e.m and representative from two independent experiments with n ≥ 4 per group. **P < 0.01, **** P < 0.0001 as calculated by two-way (**b**) or one-way (**d**) ANOVA.

### Mice deficient in type 2 immunity develop an acquired inflammatory skin phenotype

While evaluating IL-4Ra^−/−^ mice, which were maintained as homozygous animals in the SPF facility at UCSF, we noted that these mice, as well as similarly maintained mice deficient in type 2 immunity (IL-4/IL-13^−/−^, STAT6^−/−^), all acquired a progressive skin phenotype with prominent facial folds and ‘racoon’ eyes with loss of coat pigmentation beginning 2-3 months after birth that increased with age (Fig. 3a). Comparison of epidermal populations from these strains with WT C57BL/6 mice by CyTOF (Fig. 3b) and flow cytometry revealed marked ILC2 expansion, but also significant increases in CD4 and CD8 T cells, as well as Tregs (Fig. 3c,d and Supplementary Fig. 3a–e). In gut and lung, strong activation of tissue ILC2s drives proliferation and niche extrusion into the blood, allowing these cells and their cytokines to affect distal tissues^13–15^. Similarly, massive skin inflammation and ILC2 expansion in type 2 immune-deficient mice were accompanied by increases in circulating ILC2s which showed the characteristic IL-18R+ skin phenotype (Supplementary Fig. 4a,b). The increase in blood ILC2s in type 2 immune-deficient mice was associated with elevations of serum IL-5 and IL-13, but also in serum IL-22 (Supplementary Fig. 4c), consistent with studies revealing transition of activated skin ILC2s to an inflammatory Type 2/3 phenotype in both mice and humans^16, 17^.

Histologic examination of the skin from type 2 immune-deficient strains revealed epidermal acanthosis, aberrant HF morphology, and enlarged sebaceous glands (Fig. 3e). We also observed increased proliferation and number of HFSCs, and increased number of HF in these strains (Supplementary Fig. 3f–h). These observations in type 2 immune-deficient mice are consistent with the loss of negative regulation of epithelial proliferation by type 2 cytokines (Fig. 2 and Supplementary Fig. 2); recent studies showed similar impact of these cytokines on intestinal epithelium^18^. Unexpectedly, these alterations were accompanied by dramatic overgrowth of follicle-associated mites with characteristics of *Demodex* species (Fig. 3e), a common obligate ectoparasite of wild mice^19^ and a close relative of *Demodex folliculorum* in humans^20^. We used morphologic criteria of mites from hair pluck specimens (Fig. 3f) and sequencing of the 18S ribosomal RNA gene to identify the mites as *Demodex musculi* (Supplementary Fig. 5). Based on the robust chitinous exoskeleton of *Demodex*, we generated an enhanced GFP-chitin-binding domain protein (eGFP-CBP) and performed immunofluorescence (IF) of involved skin, thus highlighting the intensive mite infestation in essentially all HF and sebaceous glands of these type 2 immune-deficient mice (Fig. 3g). Additionally, we established an assay for the *Demodex* chitin synthase gene (CHS), which is highly conserved among mite species^21^, to enable a sensitive semi-quantitative PCR index for parasite skin burden in type 2-deficient mice that was absent in separately maintained WT mice (Fig. 3h).

**Fig. 3.**
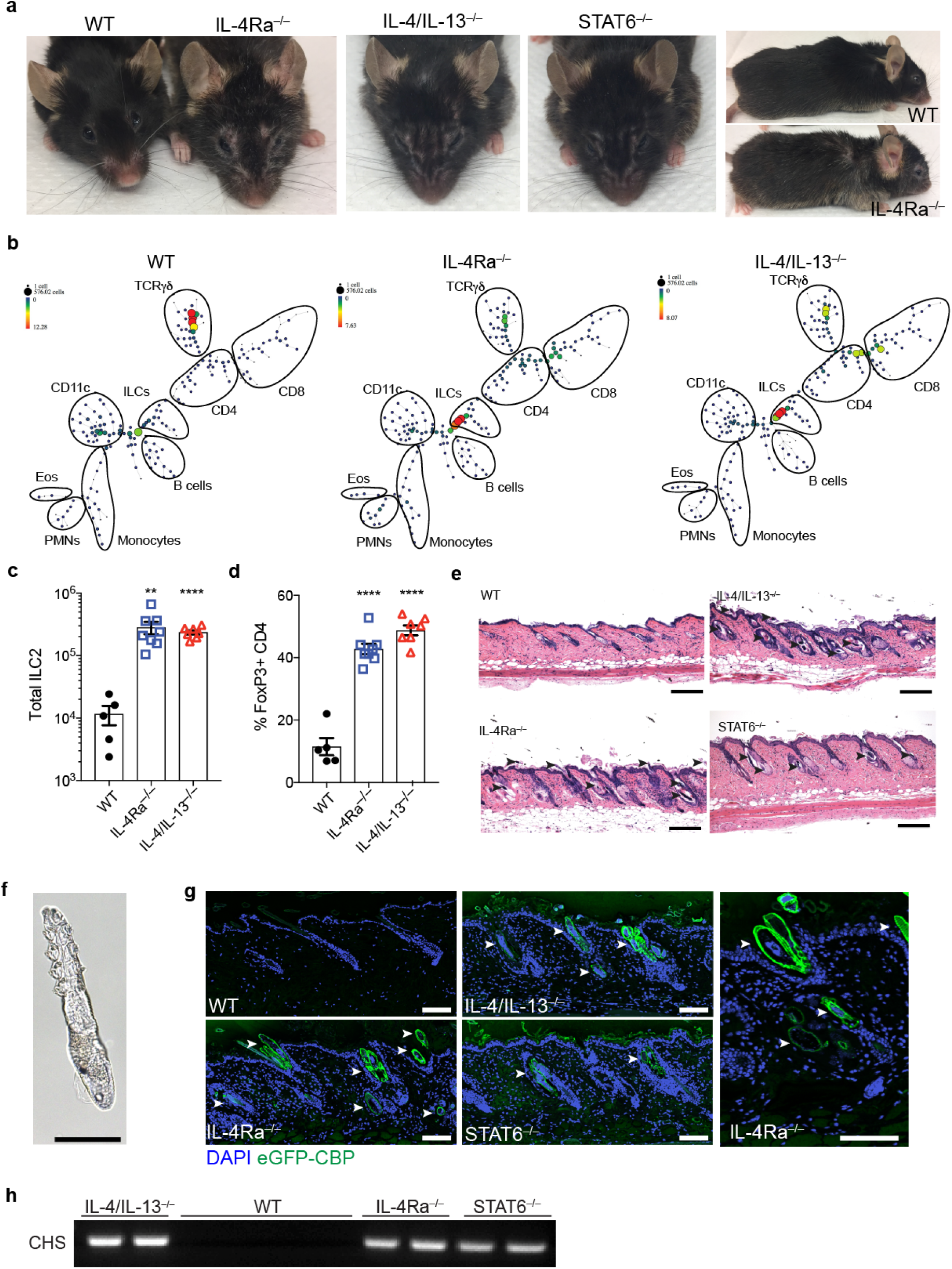
Loss of type 2 immunity is associated with an unusual skin phenotype. **a**, Representative facial and fur phenotype of wild type (WT), IL-4Ra^−/−^, IL-4/IL-13^−/−^ and STAT6^−/−^ mice at homeostasis. Mice shown are retired male breeders at 18-20 months of age. Images are representative of n ≥ 5 mice of both sexes for each genotype. **b**, SPADE representation of CD45+ cells from epidermal fraction of 8-10 weeks old WT, IL-4Ra^−/−^, and IL-4/IL-13^−/−^ mice. One representative plot shown, n = 4 individual mice per genotype. **c**, Number of ILC2s in skin from WT, IL-4Ra^−/−^, and IL-4/IL-13^−/−^ mice. **d**, Frequencies of skin Tregs (CD3+CD4+FoxP3+, as a percentage of total CD4) in skin from WT, IL-4Ra^−/−^, and IL-4/IL-13^−/−^ mice at homeostasis. **e**, Skin sections from WT, IL-4/IL-13^−/−^, IL-4Ra^−/−^ or STAT6^−/−^ mice stained with H&E. Arrowheads highlight infestation by Demodex mites in hair follicles and sebaceous glands. Scale bar, 200 μm. **f**, Brightfield image of a Demodex mite from a fur pluck of IL-4Ra^−/−^ mice. Scale bar, 50 μm. **g**, Skin sections from WT, IL-4/IL-13^−/−^, IL-4Ra^−/−^ or STAT6^−/−^ were stained for chitin (green) using an enhanced GFP chitin-binding domain fusion (eGFP-CBP) and DAPI (blue). Arrowheads highlight the Demodex mites. Scale bar, 100 μm. **h**, PCR for *Demodex* chitin synthase (CHS) from fur plucks. Data are from one representative experiment (**b**), or pooled from multiple independent experiments (**c**, **d**). * P < 0.05, ** P < 0.01, *** P < 0.001, **** P < 0.0001 by two-tailed Student’s t test (vs. WT).

### *Demodex* outgrowth in type 2 immune-deficient mice

To assess the role of mite infestation in the inflammatory dermopathy, we introduced uninfected WT mice into the colony to establish littermates for backcrossing to generate WT, heterozygous and homozygous IL-4Ra-deficient mice derived from affected animals in the same environment. Among the F2 neither WT, heterozygous, nor KO littermates developed the skin or facial phenotypes when derived from an affected homozygous KO grandparent and heterozygous parents (Supplementary Fig. 6), whereas the F1 KO mice derived from the intercross of affected IL-4Ra^−/−^ x unaffected heterozygous IL-4Ra^+/–^ mice developed the inflammatory skin phenotype (Supplementary Fig. 7a). In each case, appearance of the skin and facial phenotype was accompanied by increases in skin ILC2s, CD4 T cells and Tregs, as well as the presence of IL-13 and IL-22 in serum and an increase in follicular mites (Supplementary Fig. 7b–i), and consistent with a role for type 2 immunity, even in heterozygous genotypes, in suppressing *Demodex* outgrowth associated with inflammatory dermopathy.

Re-derived homozygous IL-4Ra-deficient mice bred in separate rooms of the SPF facility did not acquire the cutaneous phenotype, and showed no expansion of skin ILC2s; these mice were free of *Demodex* as assessed both by histology and PCR for *Demodex* chitin synthase (Fig. 4a,g,h). When housed with affected IL-4Ra-deficient mice, however, type 2 immune-deficient mice developed the characteristic skin phenotype over 2-3 months, in association with skin inflammation, aberrant HF morphology, the appearance of skin ILC2s in blood with circulating IL-13 and IL-22, and proliferation of follicular *Demodex* (Fig. 4b–f, and Supplementary Fig. 8). Histologic examination showed marked epidermal acanthosis, altered follicle morphologies and inflammatory infiltrates, which did not occur in unaffected littermates or WT mice (Fig. 4g). Affected mice also showed increases in baseline epithelial proliferation as assessed by EdU incorporation (Supplementary Figs. 3d and 9).

**Fig. 4.**
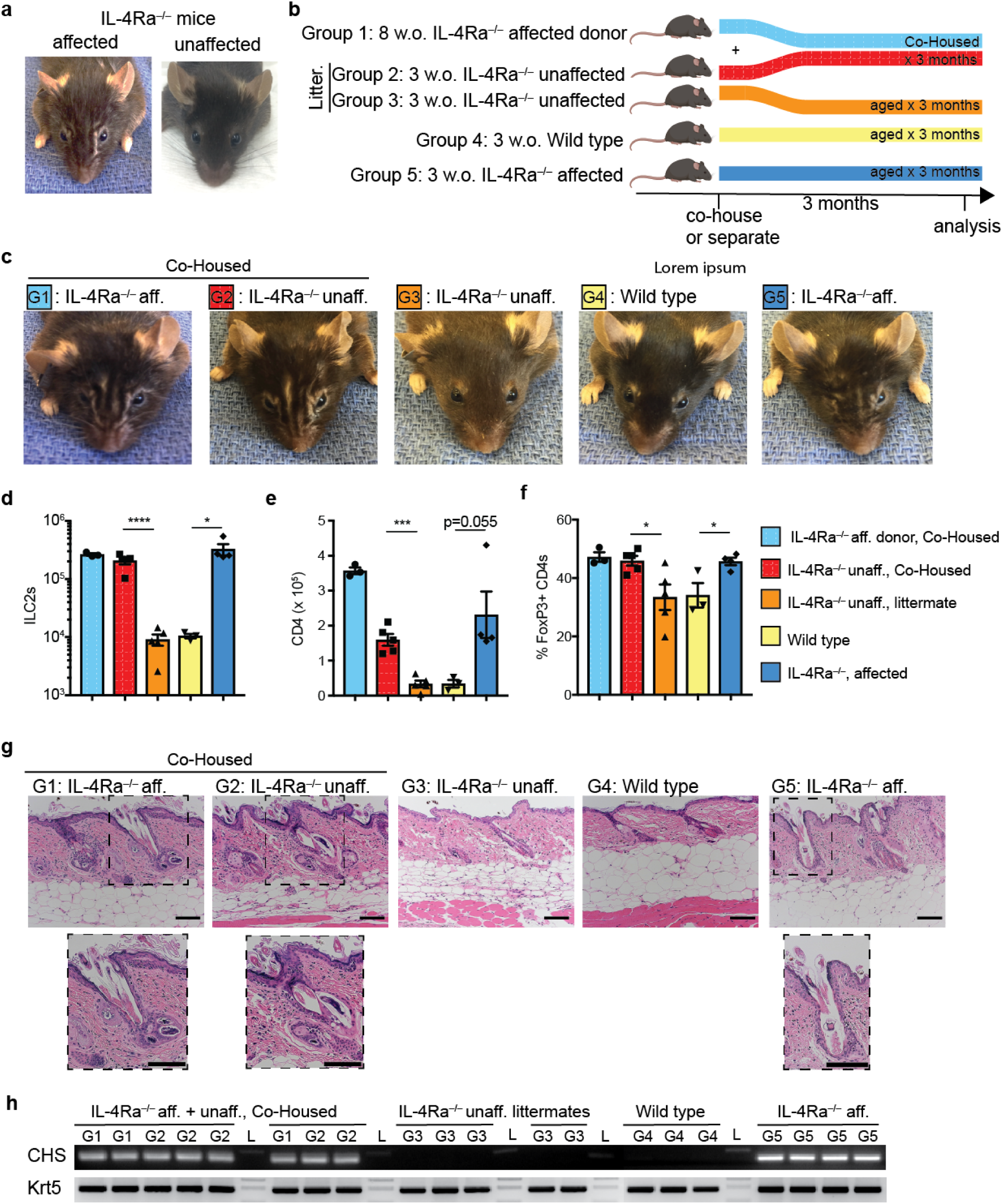
Co-housing *Demodex* infested and non-infested IL-4Ra^−/−^ mice transfers the phenotype. **a**, Facial appearance of IL-4Ra^−/−^ mice infected (affected) or uninfected (unaffected) with *Demodex musculi*. **b**, Schematic of the co-housing experiment. 8 week-old (w.o.) IL-4Ra^−/−^ mice infected with *Demodex* (Group 1) were co-housed with unaffected 3 w.o. IL-4Ra^−/−^ mice (Group 2). Unaffected littermates of co-housed mice (Group 3) were allowed to age concurrently. WT (Group 4) and IL-4Ra^−/−^ from known Demodex infected parents (Group 5) were aged as independent controls. **c**, Representative mice at 3 months after the co-housing experiment. G1-G5 indicate experimental groups as outlined in (**b**). **d,e**, Number of skin ILC2 (**d**) and CD4^+^ (**e**) cells in back skin. **f**, Frequency of Tregs (as percentage of total CD4) in back skin. **g**, Sections from back skin were stained with H&E. Inset highlights *Demodex* infestation of the hair follicle and sebaceous glands. Scale bar, 100 μm**. h**, PCR for *Demodex* chitin synthase gene (CHS, top) or genomic DNA for the keratin 5 gene (Krt5, bottom) from 2mm punches of back skin. Data are from one representative experiment of two independent experiments. Statistical significance shown by * P < 0.05, ** P < 0.01, *** P < 0.001, **** P < 0.0001 by two-tailed Student’s t test.

We next housed young IL-4Ra^−/−^ mice from affected colonies before they acquired the phenotype with WT mice to ascertain the role of type 2 immunity in limiting new infestation. While affected IL-4Ra^−/−^ mice subsequently developed the cutaneous and facial phenotype, the co-housed WT mice remained visually unaffected (Fig. 5a). Whereas WT mice were *Demodex*-free at the start of co-housing, however, these animals became infected by low numbers of mites as established by histologic exam, IF, and PCR for the *Demodex* chitin synthase gene (Fig. 5b,c and Supplementary Figs. 10, 11a). Infected WT mice developed expansion of skin ILC2s, characterized by increased IL-13 and IL-5 expression in skin but without appearance of cytokines in blood; expansion of activated CD4 T cells and Tregs was modest. Thus, tissue ILC2 activation represents a key response to this commensal ectoparasite of HF (Fig. 5d–g, and Supplementary Figs. 10, 11b). Indeed, when co-housed with affected IL-4Ra-deficient mice, unaffected Rag-deficient mice, which lack adaptive T cells and Tregs, also remained visually unaffected despite *Demodex* infection as shown by expansion and activation of skin ILC2s, and evidence for restraints on infection as assessed by histology, fewer and more superficially localized mites by immunofluorescence, and an attenuated chitin synthase PCR signal as compared to co-housed IL-4Ra-deficient mice (Fig. 5h–k, and Supplementary Fig. 11c–f).

**Fig. 5.**
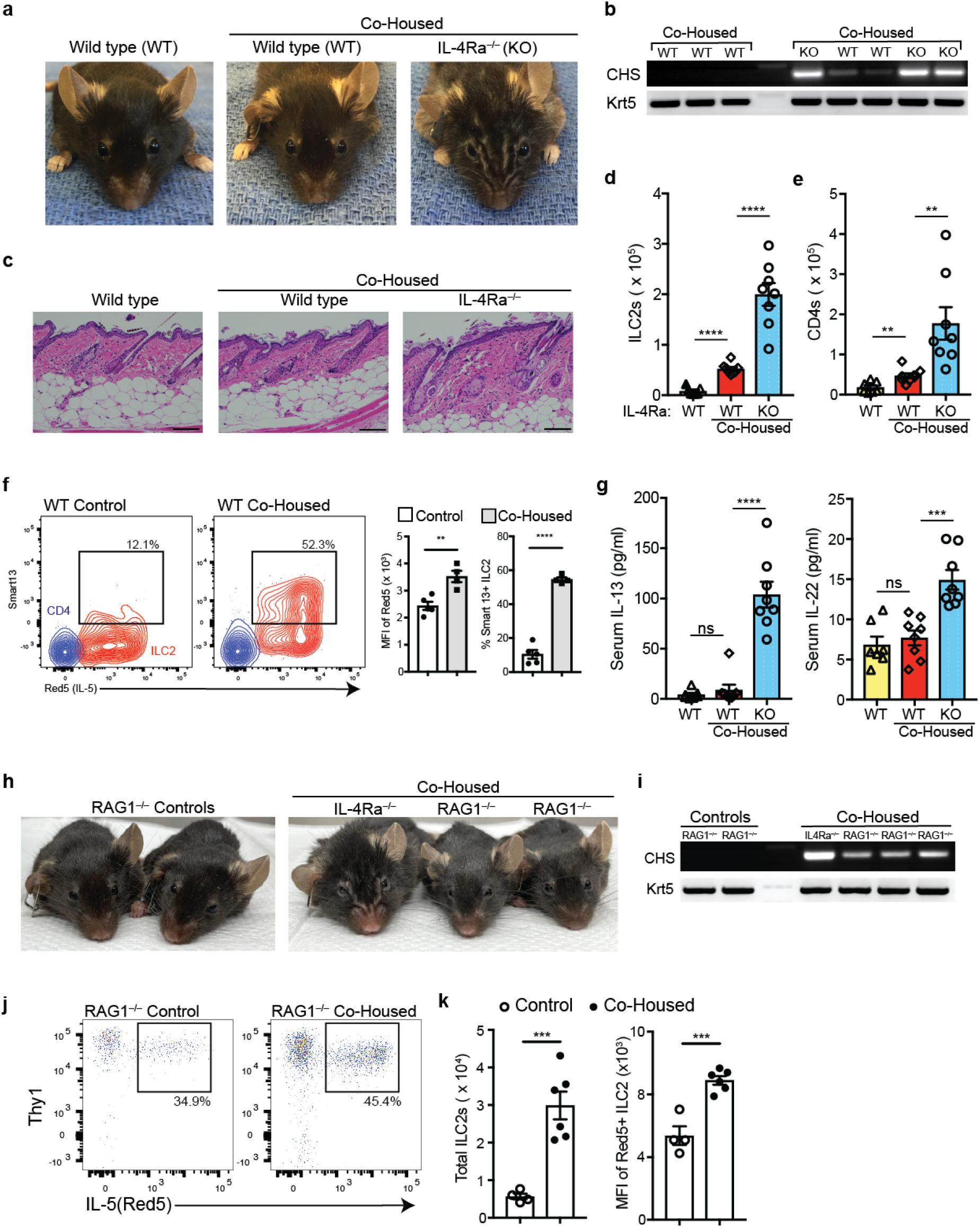
Wild type and Rag-deficient mice restrain *Demodex* infection. **a**, Representative wild type (WT), and co-housed WT with *Demodex* infested IL-4Ra^−/−^ (KO) mice. Three week old WT and infested IL-4Ra^−/−^ mice were co-housed for 12 weeks. **b**, PCR for *Demodex* chitinase synthase gene (CHS, top) or genomic DNA for the keratin 5 gene (Krt5, bottom) from back skin. **c**, H&E-stained sections of back skin from WT, or co-housed WT and IL-4Ra^−/−^ mice. **d,e**, Number of skin ILC2s (**d**) and total CD4 (**e**). **f**, Quantification of IL-5 and IL-13 expression by skin ILC2s. WT *Il5^Red5^*; *I13^Sm^*^13^ littermates were separated (Control) or co-housed with infested IL-4Ra^−/−^ mice (Co-housed). Flow cytometry plots show IL-5 and IL-13 expression by skin ILC2 (Red) or CD4 (blue). **g,** Quantification of serum IL-13 and IL-22. **h**, Representative images of Rag1^−/−^;IL-5^Red5/Red5^ littermates that were maintained by themselves (Control) or Co-Housed with Demodex infested IL-4Ra^−/−^ mice (Co-Housed) for 8 weeks. **i**, PCR for *Demodex* chitin synthase gene (CHS, top) or genomic DNA for the keratin 5 gene (Krt5, bottom) from back skin. **j**, Flow cytometry plots of Red5^+^ ILC2s (pre-gated on Live CD45^+^Lin^−^ Thy1^+^, top from Rag1^−/−^ IL-5^Red5/Red5^ (control) or Rag1^−/−^;IL-5^Red5/Red5^ that were co-housed with IL-4Ra^−/−^ mice (Co-Housed). **k,** Quantification of the number of Red5+ ILC2s (left), and mean fluorescence intensity (MFI) of Red5+ (right) ILC2s in skin. Data presented as mean ± s.e.m and pooled from two independent experiments. Statistical significance shown by * P < 0.05, ** P < 0.01, *** P < 0.001, **** P < 0.0001 by two-tailed Student’s t test. ns, not significant.

### Targeted anti-parasite therapy reverses the inflammatory dermopathy

Loss of skin ILCs leads to sebaceous gland hypertrophy and altered sebum constituents, and can affect the skin microbiota^22^. To investigate a potential microbiota contribution to the skin phenotype, we performed 16S rRNA marker gene sequencing on skin samples from affected and unaffected WT and IL-4Ra^−/−^ mice (Supplementary Fig. 12a). Although bacterial diversity was similar between phenotypes, an analysis of weighted UniFrac distance between samples showed a difference in overall community composition (Supplementary Fig. 12b, p = 0.001, PERMANOVA test). Looking to the broad characteristics of the bacteria present, we found that the fraction of obligate anaerobic bacteria was higher in unaffected mice (p = 0.028, Supplementary Fig. 12c). Despite these microbiota differences, maintaining affected IL-4Ra-deficient mice on broad spectrum antibiotics had no effect on the phenotype (Supplementary Fig. 12d). In contrast, treatment of affected littermates with topical anti-mite agents moxidectin and imidacloprid^23^, but not vehicle control (ethanol), reversed the skin phenotype in association with reductions in skin ILC2s and normalization of CD4 T cells and Tregs, as well as a marked decrease in the cutaneous mite burden (Fig. 6a–f). Treated mice also had a decrease in epithelial hyperproliferation when compared to affected IL-4Ra-deficient mice (Fig. 6g). We obtained comparable results when treating IL-4/IL-13^−/−^ mice with established *Demodex* infection (Supplementary Fig. 13).

**Fig. 6.**
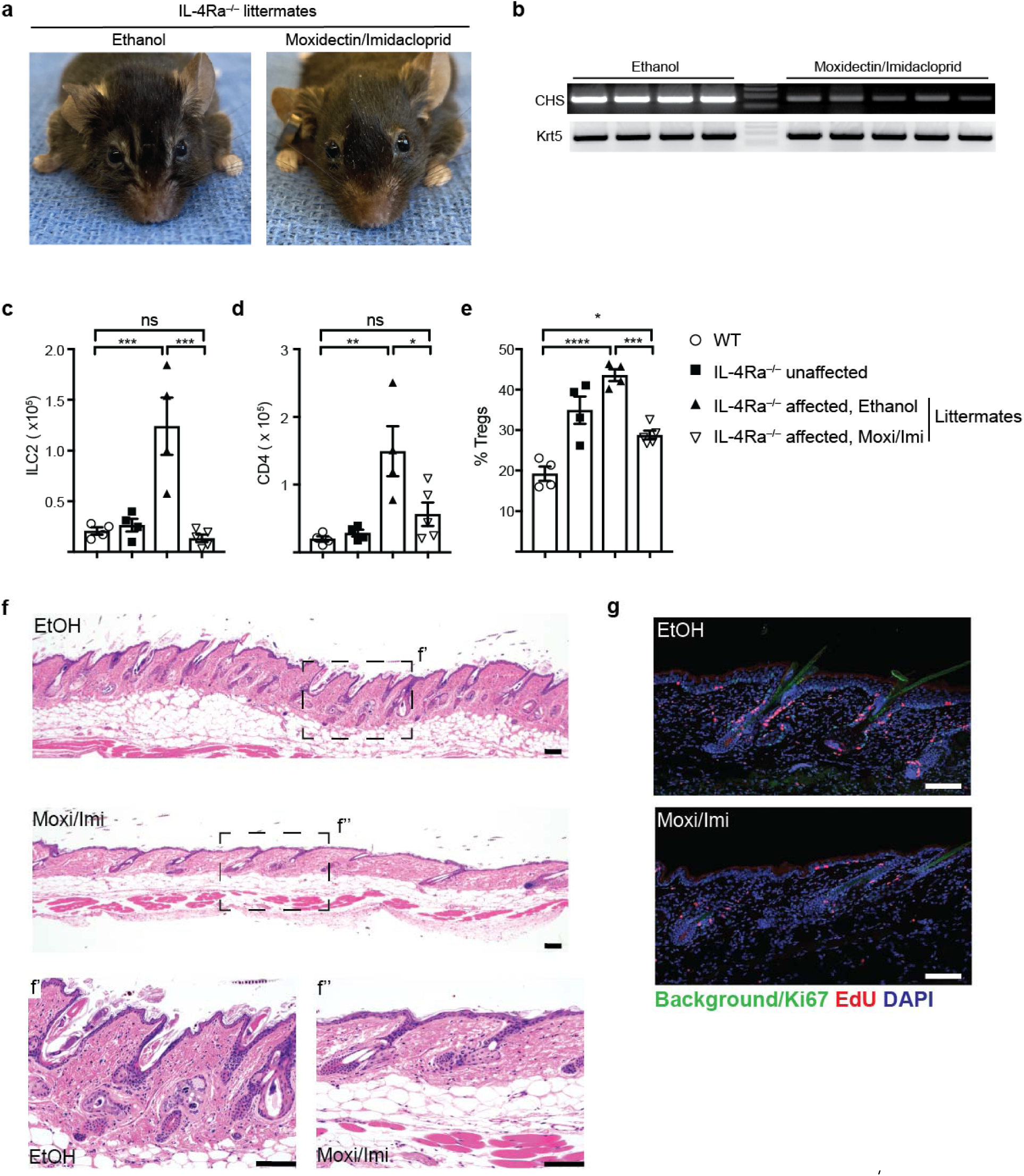
Targeted therapy reverses the phenotype in *Demodex* infested IL-4Ra^−/−^ mice. **a**, *Demodex* infested IL-4Ra^−/−^ mice littermates that were separated and treated with Ethanol (EtOH, vehicle) or Moxidectin/Imidacloprid mixture once a week for 8 weeks. **b**, PCR for *Demodex* chitin synthase gene (CHS, top) or genomic DNA for the Keratin gene (Krt5, bottom) in vehicle control (Ethanol) or Moxidectin/Imidacloprid treated mice. Each lane represents an individual mouse. **c,d**, Quantification of skin ILC2 (**c**) and CD4 (**d**) from control WT, or unaffected IL-4Ra^−/−^ or from Demodex infested IL-4Ra^−/−^ littermates that were treated with EtOH or Moxidectin/Imidacloprid (Moxi/Imi). **e**, Frequencies of Tregs in skin. **f**, Sections from back skin of control (EtOH) or moxidectin/imidacloprid (Moxi/Imi) treated mice were stained with H&E. Scale bar, 100 μm. **g**, Sections from back skin were stained for Ki67 (green), EdU (red) and DAPI (blue). Scale bar, 100 μm. Data presented as mean ± s.e.m and pooled from two independent experiments. Statistical significance shown by * P < 0.05, ** P < 0.01, *** P < 0.001, **** P < 0.0001 by one-way ANOVA. ns, not significant.

## Discussion

Outbreaks of *Demodex* mites, ubiquitous ectoparasites of feral rodents ^19, 20^, have been infrequently reported among SPF facilities, but have been noted in mouse strains deficient in type 2 immunity, as occurred here^24^. As we show by co-housing, signaling through IL-4Ra is required for limiting outgrowth of *Demodex* mites once acquired in the pilosebaceous units of the skin. The ability of Rag-deficient mice to restrain *Demodex* infestation in association with activation and expansion of skin ILC2s, focally arrayed about the hair follicles, suggests a critical role for innate type 2 immunity in control of these prevalent symbionts. Expansion of skin ILC2s was accompanied by acquisition of an ILC2/3 phenotype marked by co-expression of IL-13 and IL-22, in agreement with natural trajectories of these resident immune cells during perturbation^17, 25^, and mimicking trajectories of skin microbiota-specific adaptive CD8 T cells^26^. As shown here, IL-13 attenuates proliferation of epithelia throughout the pilosebaceous unit, and perhaps better integrates with the effects of IL-22 on epithelial differentiation^27^ to stabilize synchronous follicular development during infection that otherwise leads to aberrant follicles that allow overgrowth of the resident mites. Intriguingly, ILC2s in intestinal mucosa regulate homeostasis with the protozoan, *Tritrichomonas*^28, 29^, suggesting a role for these innate immune cells and type 2 immunity in sustaining tissue functionality in the setting of complex barrier-associated eukaryotic pathobionts. Inhibitory effects of IL-4/13 on intestinal epithelial proliferation and differentiation have been described^18^, and further work is needed to clarify these pathways. Although we cannot exclude a role for microbiota-mediated inflammation contributing to the inflammatory phenotype caused by *Demodex* outgrowth, the infection was controlled by topical therapeutics targeting the mite but not by antibiotics.

*Demodex* species are obligate symbiont ectoparasites that inhabit the HF and sebaceous glands of most mammals over millions of years of co-evolution^30, 31^. Among humans, prevalence of asymptomatic *Demodex* infection, particularly on the face, reaches essentially 100% with age, but outgrowths can be provoked by immunodeficiency and malnutrition, and have been associated with rosacea, although causality is uncertain^32, 33^. While relatively short-lived, the organisms eat sebum and keratinocyte debris and produce 1-2 dozen eggs that differentiate through intermediate larvae to generate progeny that spread to nearby follicles^34^. Mechanisms that restrain the fecundity, growth and/or maintenance of follicular mites are little studied, and the models we describe will be useful to probe these pathways as well as implications for disease in the colonized host, such as tolerance to allergens or proclivity to AD. Of note, patients treated with anti-IL4Ra antagonist for atopic dermatitis have increased incidence of conjunctivitis associated with demidocosis^35^, suggesting pathways we uncover here likely participate in control of follicular mites in humans.

## Materials and Methods

### Mice

Wild-type (C57BL/6J; Stock 000664) mice were purchased from Jackson Laboratories. *Il5*^Red5^, *Arg1*^Yarg^, and *Il13*^Smart^ reporter alleles on C57BL/6J (B6) backgrounds were bred and maintained as described^7, 36^. For some experiments, mice encoding the *Arg1*^Yarg^ and/or *Il13*^Smart^ reporter alleles that do not impact the endogenous gene expression were used as wild-type controls. *Il4ra^−/−^* (BALB/c-Il4ratm1Sz/J; 003514) and *Stat6*^−/−^ (B6.129S2(C)-Stat6tm1Gru/J; 005977) mice were initially purchased from Jackson Laboratories and backcrossed to C57BL/6J for at least eight generations as described^37^. Additional wild-type C57BL/6J and IL-4Ra^−/−^ mice on the C57BL/6J background were obtained from the UCSF colony of M.S. Fassett and K.M. Ansel. *Il4/Il13^−/−^* mice were generated as previously described and backcrossed C57BL/6J for at least eight generations^38^. Rag1^−/−^ mice (Stock 002216) were purchased from Jackson laboratory, and crossed to *Arg1*^Yarg^ and *Il5*^Red5^ reported alleles. For all experiments, sex-matched mice aged 7-14 weeks were used unless otherwise indicated. Mice were maintained under specific pathogen-free conditions. For *in vivo* cytokine treatments, mice were injected subcutaneously with IL-4 complex (IL-4c) comprised of rmIL-4 (2 μg, R&D systems) combined with anti-IL-4 antibody 10 μg, Clone Mab 11B11, BioXcell), or rmIL-13 (2 μg, R&D systems) in calcium- and magnesium-free phosphate buffered saline (PBS). For co-housing experiments, mice of the appropriate genotypes were co-housed for 8-12 weeks in the ratios noted in the figure legends. For treatment with topical anti-parasitic agents, a stock solution in 70% ethanol of moxidectin (Sigma 33746) and imidacloprid (Sigma 37894) was prepared and diluted to a working solution of (0.83 mg/ml moxidectin and 3.33 mg/ml imidacloprid) and applied topically to intrascapular skin at 3.3 mg/kg moxidectin and 13.3 mg/kg imidacloprid. Control mice were treated with 70% ethanol. For broad-spectrum antibiotics treatment, mice received drinking water supplemented with ampicillin, kanamycin, and neomycin (all 1 mg/ml), vancomycin (0.5 mg/ml), and metronidazole (2.5 mg/ml) in 1% sucrose. For EdU experiments, mice received a peritoneal injection of 1 μg of EdU (Sigma 900584) 14-18 hours prior to harvesting. All animal procedures were approved by the UCSF Institutional Animal Care and Use Committee.

### Depilation Experiment

Mice were anesthetized with isoflurane inhalation. Under anesthesia, the dorsal surface (back) hair of the animals was shaved down to the level of skin with an electric razor. A thin coat (1 fingertip unit) of Nair depilatory cream (Nivea) was applied to the shaved region for a period of 30 s before wiping clean. For monitoring of clinical hair regrowth, standardized pictures were taken with a ruler on the day of depilation (day 0) and then at days 4, 7, 9, 11, 14. Anagen induction was quantified using intensity analysis on ImageJ software (NIH, USA) at each time point or as a percent of pigmented dorsal skin relative to baseline (day 0). Tissues were analyzed for flow cytometry, stem cell proliferation, and qPCR as described below.

### Flow Cytometry and Cell Sorting

Whole skin single cell suspensions were prepared as described previously^7^. Briefly, back tissue was minced in RPMI-1640 with 5% FBS, then transferred to C tubes (Miltenyi Biotec), containing 5 ml of RPMI-1640 (Sigma R8758) supplemented with Liberase TL (0.25 mg/ml, Sigma 5401020001), and DNAse I (0.1 mg/ml; Sigma 10104159001). Samples were shaken at 250 rpm for 2 hours at 37 °C, then dispersed using an automated tissue dissociator (GentleMACS; Miltenyi Biotec) running program C. Single-cell suspensions were passed through a 70 μm filter and washed twice with RPMI containing 5% FBS. The following antibodies, all from BioLegend (unless otherwise specified, see **Supplementary Table 1**) were used at 1:300 dilution unless noted: anti-CD3 (17A2, diluted 1:200), anti-CD4 (RM4-5, diluted 1:100), anti-CD5 (53-7.3), anti-CD8α (53-6.7, diluted 1:100), anti-CD11b (M1/70), anti-CD11c (N418), anti-CD19 (6D5), anti-CD25 (PC61, diluted 1:100), anti-CD45 (30F-11, BD Biosciences), anti-CD49b (DX5; eBiosciences), anti-CD127 (A7R34, diluted 1:100), anti-CD218 (P3TUNYA, eBiosciences), anti-F4/80 (BM8), anti-Gr-1 (RB6-8C5), anti-NK1.1 (PK136), anti-NKp46 (29A1.4), anti-Thy1.2 (30-H12; diluted 1:1000); anti-human CD4 (RPA-T4, diluted 1:20; eBiosciences), anti-GATA3 (TWAJ, diluted 1:25), anti-FoxP3 (FJK-16s, Invitrogen, diluted 1:100), anti-T1/ST2 (U29-93; BD Biosciences, diluted 1:200). Live/dead cell exclusion was performed with DAPI (4′-phenylindole dihydrochloride; Roche) or LIVE/dead fixable dye (Thermofisher). Cell counts were performed using flow cytometry counting beads (CountBright Absolute; Life Technologies) per manufacturers instructions. To assay cell proliferation, the Click-IT EdU flow cytometry or imaging kits (ThermoFisher) were used according to the kit instructions. Sample data were acquired with a 5-laser LSRFortessa X-20 flow cytometer and BD FACSDiva software (BD Biosciences) and analyzed using FlowJo software (Tree Star v10.6.1).

ILC2s were sorted from reporter mice as live (DAPI–), Lin(CD3, CD4, CD8, CD11b, CD11c, CD19, NK1.1, NKp46, Gr-1, F4/80, Ter119, DX5)–CD45+Red5+ using a MoFlo XDP (Beckman Coulter). Epithelial cell preparations for flow cytometry or CyTOF analyses were prepared as previously described^10, 39^. Briefly, mice were shaved and dorsal skin was harvested and trimmed of fat. The tissue was floated in 4 ml of 0.25% Trypsin-EDTA (ThermoFisher) and incubated for 90 min at 37 °C. The epidermis was scrapped from the dermis and mechanically separated by pipetting into complete media (RPMI-1640 supplemented with 10% FBS, HEPES, Penicillin/Streptomycin). The cell suspension was passed through a 100 μm strainer and washed with 10 ml of complete media. The cell suspension was then pelleted, filtered through a 40 μm filter and aliquoted for cell staining. The following antibodies were used for staining epithelial fractions: anti-Sca1 (D7, 1:300); anti-MHCII (M5/114.15.2, 1:400); anti-CD326 (G8.8, 1:300); anti-CD45 (30F-11, 1:400); anti-CD34 (RAM34, 1:200); anti-CD49f (GoH3, 1:300).

### Mass Cytometry

#### Sample preparation

Mass cytometry was performed as described^40^. Briefly, single cell suspension from the epidermal fraction of the various mice strains were prepared. To test the responsiveness to type 2 immune signaling, mice were treated with s.c. IL-4 complex for 30-60 min prior to harvest. IdU (200 μl of 5mM stock solution, Sigma I7125) was injected 20-30 min prior to harvesting. Cells were incubated with 25 μM Cisplatin for 1 min and fixed with paraformaldehyde at 1.6% for 10 min at room temperature (RT). Cell pellets were stored at –80 °C until analysis. Prior to antibody staining, cells were thawed and mass tag cellular barcoding of prepared samples was performed by incubating cells with distinct combinations of isotopically-purified palladium ions chelated by isothiocyanobenzyl-EDTA in 0.02% saponin in PBS as previously described^41, 42^. After two washes with staining media (PBS + 0.5% BSA + 0.02% NaN_3_), sets of 20 barcoded samples were pooled together and washed once more with staining media. Antibody staining was performed using a panel of antibodies and metals that were purchased directly conjugated (Fluidigm) or conjugated using 100 μg of antibody lots combined with the MaxPAR antibody conjugation kit (Fluidigm) according to the manufacturer’s instructions and detailed in **Supplementary Table 2**. Surface marker antibodies were added to a 500 μL final reaction volumes and stained for 30 min at RT on a shaker. Following staining, cells were washed twice with PBS with 0.5% BSA and 0.02% NaN3. to remove remaining methanol. Then cells were permeabilized with 4 °C methanol for 10 min at 4 °C followed by two washes in PBS with 0.5% BSA and 0.02% NaN_3_ to remove remaining methanol. The intracellular antibodies were added in 500 μ L for 30 min at RT on a shaker. Cells were washed twice in PBS with 0.5% BSA and 0.02% NaN_3_ and stained with 1 mL of 1:4000 191/193Ir DNA intercalator (Fluidigm) diluted in PBS with 1.6% PFA overnight. Cells were washed once in PBS with 0.5% BSA and 0.02% NaN_3_ and twice with double-deionized (dd)H_2_0. We analyzed 1 × 10^6^ cells per animal, per tissue, per condition consistent with generally accepted practices in the field.

#### Bead Standard Data Normalization

Just before analysis, the stained and intercalated cell pellet was resuspended in ddH_2_O containing the bead standard at a concentration ranging between 1 and 2 x 10^4^ beads/ml as described^43^. The bead standards were prepared immediately before analysis, and the mixture of beads and cells was filtered through filter cap FACS tubes (BD Biosciences) before analysis. Mass cytometry files were normalized together using the mass cytometry data normalization algorithm^43^, which uses the intensity values of a sliding window of these bead standards to correct for instrument fluctuations over time and between samples.

#### Scaffold Map Generation

Total live leukocytes (excluding erythrocytes) were used for all analyses. Cells from each tissue for all animals were clustered together (rather than performing CLARA clustering on each file individually as originally implemented in^44^.) Cells were deconvolved into their respective samples. Cluster frequencies or the Boolean expression of Ki67 for each cluster were passed into the Significance Across Microarrays algorithm^45, 46^, and results were tabulated into the Scaffold map files for visualization through the graphical user interface. Cluster frequencies were calculated as a percent of total live nucleated cells (excluding erythrocytes). Scaffold maps were then generated as previously reported^44, 47^. All analyses were performed using the open source Scaffold maps R package available at http://www.github.com/spitzerlab/statisticalScaffold.

#### Spade and viSNE Generation

viSNE and Spade analyses were performed with Cytobank (www.cytobank.org).

### Cytokine quantification

Cytokine levels in mouse serum were measured according to the manufacturer’s protocol using the following Cytokine Bead Array Flex Sets (BD): IL-4 (#558298), IL-5 (#558302), IL-13 (#558349), and IL-17A (#560283). The data were analyzed using Flow Cytometric Analysis Program (FCAP) Array software (BD). Serum IL-22 was measured using LEGEND MAX^TM^ Mouse IL-22 ELISA Kit (BioLegend, #436307) and analyzed using a SpectraMax M2 microplate reader using SoftMax Pro software (Molecular Devices).

### RNA preparation and qRT-PCR

ILC2s were sorted into RLT Plus lysis buffer (Qiagen) and stored at – 80 °C until processing using RNeasy Micro Plus kit (Qiagen) per manufacturer’s protocol. For quantitative reverse transcriptase PCR analyses, RNA was reverse transcribed using SuperScript VILO cDNA synthesis kit (ThermoFisher) per the manufacturer’s instructions. The cDNA was used to the targets of interest using Power SYBR Green PCR master mix (ThermoFisher) in a StepOnePlus cycler (Applied Biosystems). The following primers sets from PrimerBank were used: *Il13:* Il13-Forward 5’-CCTGGCTCTTGCTTGCCTT-3’ and Il13-Reverse, 5’-GGTCTTGTGTGATGTTGCTCA-3’; *Rps17*: Rps17-Forward 5’-CCAAGACCGTGAAGAAGGCTG-3’, Rps17-Reverse 5’-GCTGGGGATAATGGCGATCT-3’. Gene expression was normalized to *Rps17* using the ΔCt method.

### Chitin Synthase and *Demodex* 18S rRNA gene PCR

For analyses of demodex burden in skin, three random 2 mm punch biopsies of back caudal skin were digested in digest buffer (10 mM Tris pH 8.0, 100 mM NaCl, 10 mM EDTA pH 8.0, 0.5% SDS and 200 μg/ml Proteinase K (Thomas Scientific)) and the genomic DNA was resuspended in 30 μl of TE. The following primers and PCR conditions were used to detect the highly conserved *Demodex* spp. chitin synthase (CHS) gene (GenBank No. AB080667): CHS-Forward 5’-GAAGCGGCGAGTAATGTTCATC-3’; CHS-Reverse 5’-CCTGACTCCATCTTTTACGATGTC-3’; Cycling conditions 95 °C 3’, (95 °C 30”, 52 °C 30”, 72 °C 30”) x 30 cycles, 72 °C 7’. Amplification of the murine Keratin 5 gene was used as a loading control using the following primers and PCR conditions: Keratin5-Forward: 5’-TCTGCCATCACCCCATCTGT-3’; Keratin5-Reverse: 5’-CCTCCGCCAGAACTGTAGGA-3’, 95 °C 3’, (95 °C 30”, 60 °C 30”, 72 °C 30”) x 30 cycles, 72 °C 7’. PCR products were separated by a 2.5% agarose gel with ethidium bromide and visualized by a GelDoc (BioRad). *Demodex musculi* was confirmed by first amplifying the 18S rRNA gene from DNA samples purified from the hair scrapes alone of affected mice using the universal 18S primers and PCR conditions: 18S-Forward 5’-TCCAAGGAAGGCAGCAGGCA-3’; 18S-Reverse 5’-CGCGGTAGTTCGTCTTGCGACG-3’, 95 °C 3’, (95 °C 30”, 58 °C 30”, 72 °C 30”) x 30 cycles, 72 °C 7’. PCR amplicons (532 bp) were purified and subsequently cloned into pCR4-TOPO plasmid by TOPO-TA cloning (Thermo Fisher) for sequencing. Putative *Demodex* spp. sequences from individual mouse (>3) were identical to the *Demodex musculi* 18S rRNA gene (GenBank No. JF834894).

### Preparation of eGFP-CBP for immunofluorescence

Enhanced GFP was cloned into pGV358 (gift from Kushol Gupta, Univ. of Pennsylvania), a pETDuet based vector that has an Mxe intein in frame at the C-terminus followed by the non-cleavable hexahistidine tagged chitin-binding domain from *Bacillus circulans* WL-12 Chitinase A1 (eGFP-CBP). Construct expressing eGFP-CBP (gift from Sy Redding, Univ. of California, San Francisco) was transformed into BL21(DE3) cells and expressed overnight in 50 mL LB media at 37 °C. 10 mL of overnight culture was diluted in 1L LB media and expressed at 37 °C until OD_600_ = 1-1.2. Culture was induced with 1 mL 1M IPTG, and expression continued overnight at 19 °C. We added protease inhibitor (Roche #11836170001) at the temperature change to minimize proteolysis of the expressed protein. Cells were centrifuged at 4,000 x g at 4 °C for 20 mins. The pelleted cells were resuspended in 50 mL of lysis buffer (100 mM Tris pH 8.5, 150 mM NaCl, 30 mg Lysozyme, protease inhibitor, 1 µL universal nuclease (ThermoFisher Scientific #88700)) per 1 L of culture, then lysed using sonication for 6 mins at 50% power. Lysed cells were centrifuged at 10,000 x g at 4 °C for 15-30 mins. The lysate supernatant was filtered with a 0.22 µm filter. The filtered lysate was bound to a 5 mL HisTrap FF column (Cytiva #17525501), washed with 100 mM Tris pH 8.5, 150 mM NaCl, 50 mM imidazole pH 7.0, and eluted with a gradient into 100 mM Tris pH 8.5, 150 mM NaCl, 500 mM imidazole pH 7.0. Fractions were selected based on presence of molecular weight band at 56.8 kDa, corresponding with eGFP-CBP, as assessed via SDS-PAGE. Fractions were pooled and dialysed overnight into 100 mM Tris pH 8.5, 150 mM NaCl, 5% glycerol w/v. The protein solution was concentrated and separated via size-exclusion chromatography on a HiLoad 16/600 Superdex 75 pg (Cytiva #28989333). Fractions were selected based on purity as assessed via SDS-PAGE.

### Fixed tissue preparation and immunofluorescence staining

For immunohistochemistry, 3-4 mm wide cephalad-caudal strips of back skin were cut and fixed in 2% paraformaldehyde for 2-6 hrs at room temperature, followed by 2 washes in D-PBS, and incubation in 30% (w/v) sucrose in D-PBS at 4 °C overnight. Portions of the tissue sections were submitted for routine paraffin embedding and hematoxylin and eosin staining using UCSF Histology and Biomarker Core. For frozen sections, tissues were embedded in Optimal Cutting Temperature Compound (Tissue-Tek) and stored at –80 °C until further processing. Tissue sections (8-12 μm) were cut on a Cryostat (Leica). Immunohistochemistry was performed in Tris/NaCl blocking buffer (0.1□M Tris-HCl, 0.15□M NaCl, 5□μg/ml blocking reagent (Perkin Elmer), pH 7.5) as follows at RT: 1□hour in 5% goat serum, 1-2□hours in primary antibody, 60□min secondary antibody (Goat anti-Rabbit AF555, 1:1000), 5□min of a 1:1000 dilution of DAPI (Stock 5 mg/ml in PBS), Roche) and mounted in Mounting Media (Vector Labs). The following primary antibodies were used in this study: Living colors DsRed Polyclonal antibody (Rabbit, 1:300, Takara Bio #632496); FITC Anti-mouse Ki67 (1:100, Invitrogen #11-5698-82), and FITC Mouse Anti-E-Cadherin (1:400, BD Biosciences #612131). For detection of chitin in skin tissue sections, 4-5 μm thick paraffin-embedded sections were baked at 60 °C for 30 min, treated with xylenes, and washed with 100% ethanol. The sections were incubated with an eGFP-CBP fusion protein (5 μg/ml) and DAPI (5 μg/ml) in PBS at RT for 60 min, washed 3 times with PBS and mounted using ProLong Gold antifade (Thermo Fisher, P36930). Images were acquired using a Zeiss immunofluorescence AxioImager M2 upright microscope running AxioVision software (Zeiss) or an inverted Nikon A1R confocal microscope running NIS Elements software (Nikon).

### 16S rRNA gene sequence analysis

#### Sample collection

Mice facial and back skin were swabbed at the end of the experiment using a sterile foam tipped applicator (Puritan) that was moistened with yeast cell lysis buffer (Lucigen) and collected into sterile 2 ml tubes (Eppendorf). All samples were immediately snap frozen in dry ice and stored at –80 °C until DNA extraction.

#### DNA extraction and V1V3 Library Prep

DNA was extracted from samples using the DNEasy PowerSoil Pro Kit (Qiagen) according to the manufacturer’s protocol. Barcoded PCR primers annealing to the V1-V3 region of the 16S rRNA gene (Fwd, 5’-AGAGTTTGATCCTGGCTCAG-3’; Rev, 5-ATTACCGCGGCTGCTGG-3’) were used for library generation. PCR reactions were carried out in quadruplicate using Q5 High-Fidelity DNA Polymerase (NEB, Ipswich, MA). Each PCR reaction contains 0.5 uM of each primer, 0.34 U Q5 Pol, 1X Buffer, 0.2 mM dNTPs, and 5.0 ul DNA in a total volume of 25 ul. Cycling conditions are as follows: 1 cycle of 98 C for 1 min; 30 cycles of 98 C for 10 sec, 56 C for 20 sec, and 72 C for 20 sec; 1 cycle of 72 C for 8 min. After amplification, quadruplicate PCR reactions were pooled and then purified using a 1:1 volume of SPRI beads. DNA in each sample was quantified using PicoGreen and pooled in equal molar amounts. The resulting library was sequenced on the Illumina MiSeq using 2×300 bp chemistry. Extraction blanks and DNA free water were subjected to the same amplification and purification procedure to allow for empirical assessment of environmental and reagent contamination.

#### Bioinformatics processing

Sequence data were processed using QIIME2^48^. Read pairs were processed to identify amplicon sequence variants with DADA2^49^. Taxonomic assignments were generated by comparison to the Greengenes reference database^50^, using the naive Bayes classifier implemented in scikit-bio^51^. A phylogenetic tree was inferred from the sequence data using MAFFT^52^.

#### Statistical analysis for microbiome data

Data files from QIIME were analyzed in the R environment for statistical computing, using the QIIMER library, which we developed (http://cran.r-project.org/web/packages/qiimer). Global differences in bacterial community composition were visualized using Principal Coordinates Analysis of weighted and unweighted UniFrac distance^53, 54^. Community-level differences between sample groups were assessed using the PERMANOVA test, which allows sample-sample distances to be applied to an ANOVA-like framework^55^. Linear mixed-effects models were used to ascertain differences in alpha diversity and taxonomic abundances.

### Statistical analysis

All experiments were performed using randomly assigned mice without investigator blinding. All data points and n values reflect biological (individual mice) replicates. Experiments were pooled whenever possible, and all data were analyzed using Prism 7 (GraphPad Software) by comparison of means using unpaired two-tailed Student’s *t* tests or one-way ANOVA for multiple comparison as indicated in the figure legends. Data in all figures displayed as mean ± s.e.m. unless otherwise indicated.

## Acknowledgments

We thank M. Ji and M. Conseco for technical expertise and mouse colony maintenance, Z. Wang for cell sorting, J. Bolen and X.Y. Jiang and the UCSF Biomarkers and Histology Core for assistance with histology. We thank members of the Locksley laboratory and Michael D. Rosenblum and Anthony L. DeFranco for comments on the manuscript.

## Author contributions

R.R.R.G. conceived the study, designed and performed experiments, analyzed and interpreted the data, and wrote the manuscript. M.E.K., H.-E.L., and J.L. participated in performing experiments, provided intellectual expertise, and/or helped to interpret experimental results. I.L. and D.M. performed CyTOF analyses on skin samples under the supervision and guidance of M.H.S. M.S.F. and K.M.A. provided mice for co-housing experiments. S.G.D. and K.B. contributed to 16S sequencing analysis. R.E.D. and J.S.F. designed, expressed, and purified the eGFP-CBP fusion protein. R.M.L. directed the studies and wrote the paper with R.R.R.G.

## Data and materials availability

Reagents are available upon signing a materials transfer agreement. All other data needed to evaluate the conclusions in this paper are present either in the main text or the supplementary materials.

## Supplementary Figures

**Supplementary Fig. 1.**
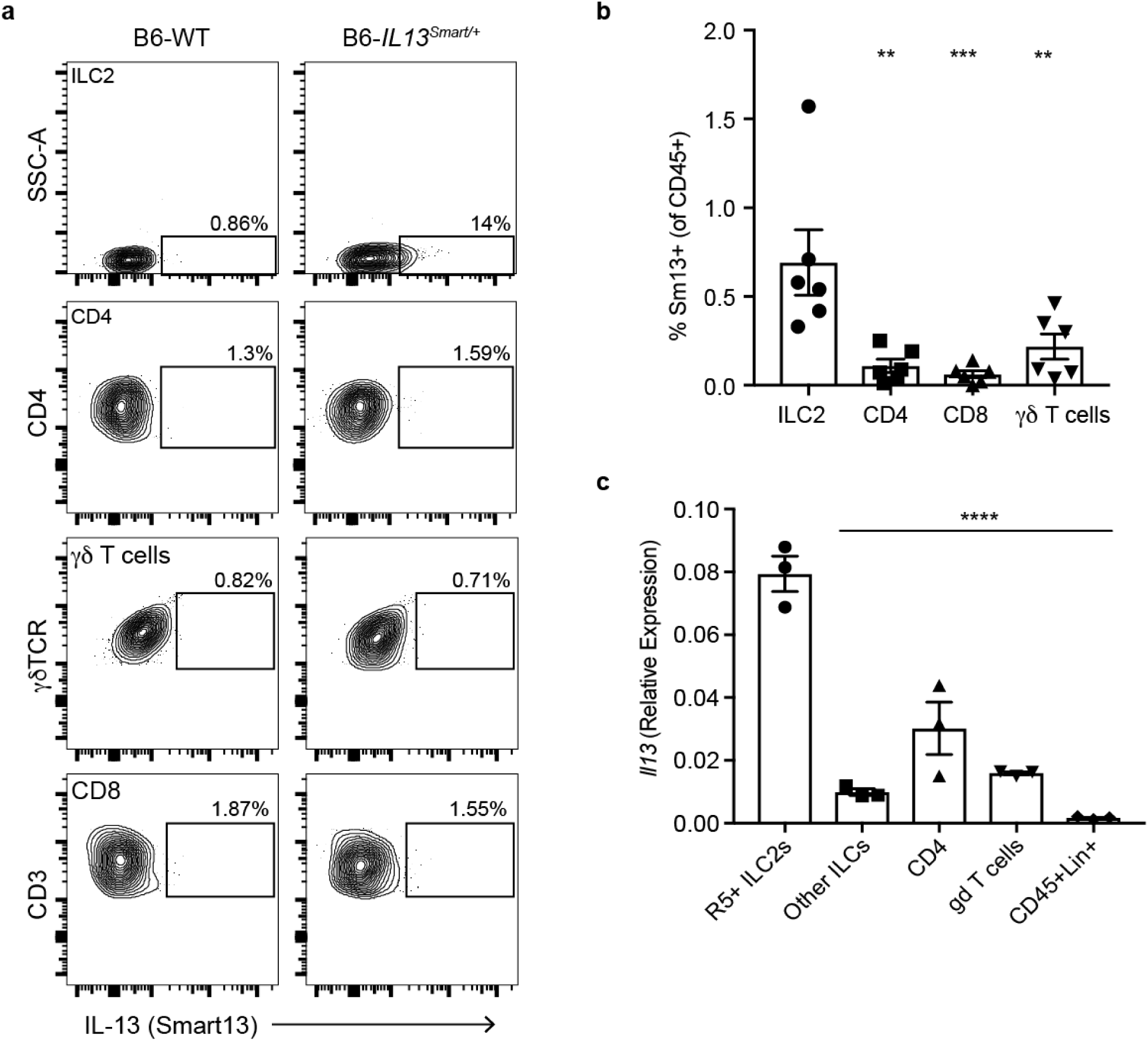
ILCs are the predominant source of IL-13 in skin in homeostasis. **a,b,** Flow cytometry plots (**a**) and relative frequency (**b**) of IL-13 expression as assessed by Smart13 (huCD4) in various lymphoid populations of adult B6-IL13^Smart13/+^ mice in homeostasis. **c**, Expression of *Il13* by qPCR in tissue resident leukocytes in the skin. ILC2s (CD45+Lin–Thy1+Red5+), other ILCs (CD45+Lin–Thy1+Red5–), CD4 (CD45+CD3+CD4+), gdT cell (CD45+CD3+gdTCR+) or the remaining leukocytes (CD45+Lin+) were sorted from 8-10 week old mice. Data are representative from two independent experiments with at least 3 individual mice per group. Statistical significance shown by ** P < 0.01, *** P < 0.001, **** P < 0.0001 by ANOVA.

**Supplementary Fig. 2.**
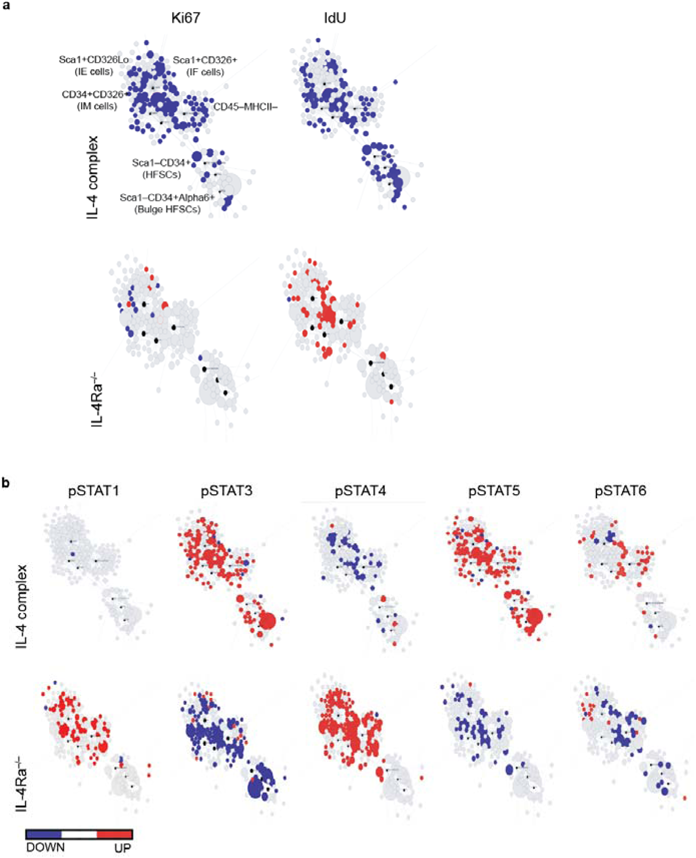
Type 2 immunity restrains cell proliferation and aberrant signaling in the epidermis. **a**, Scaffold maps of Ki67 and IdU for the CD45–epidermal cell fraction as analyzed by CyTOF (see methods). Black nodes represent canonical cell populations identified manually as noted in the top left scaffold map. Blue denotes the population has significant lower expression frequency of the marker (Ki67, or IdU, as denoted in the columns) in the experimental arm (IL-4 complex treated WT mice, top; or IL-4Ra^−/−^ mice, bottom) vs. WT controls; red denotes significantly higher frequency. n = 4 individual mice per group. **b**, Scaffold maps for phospho(p)STAT1, pSTAT3, pSTAT4, pSTAT5, or pSTAT6 in IL-4 complex treated (top) or IL-4Ra^−/−^ (bottom) mice compared to WT controls. Blue and red colors denote statistically significant decrease or increase in expression frequency, respectively, in each cluster/node compared to the control group (q < 0.05 by SAM). Data are representative n = 4 individual mice per group.

**Supplementary Fig. 3.**
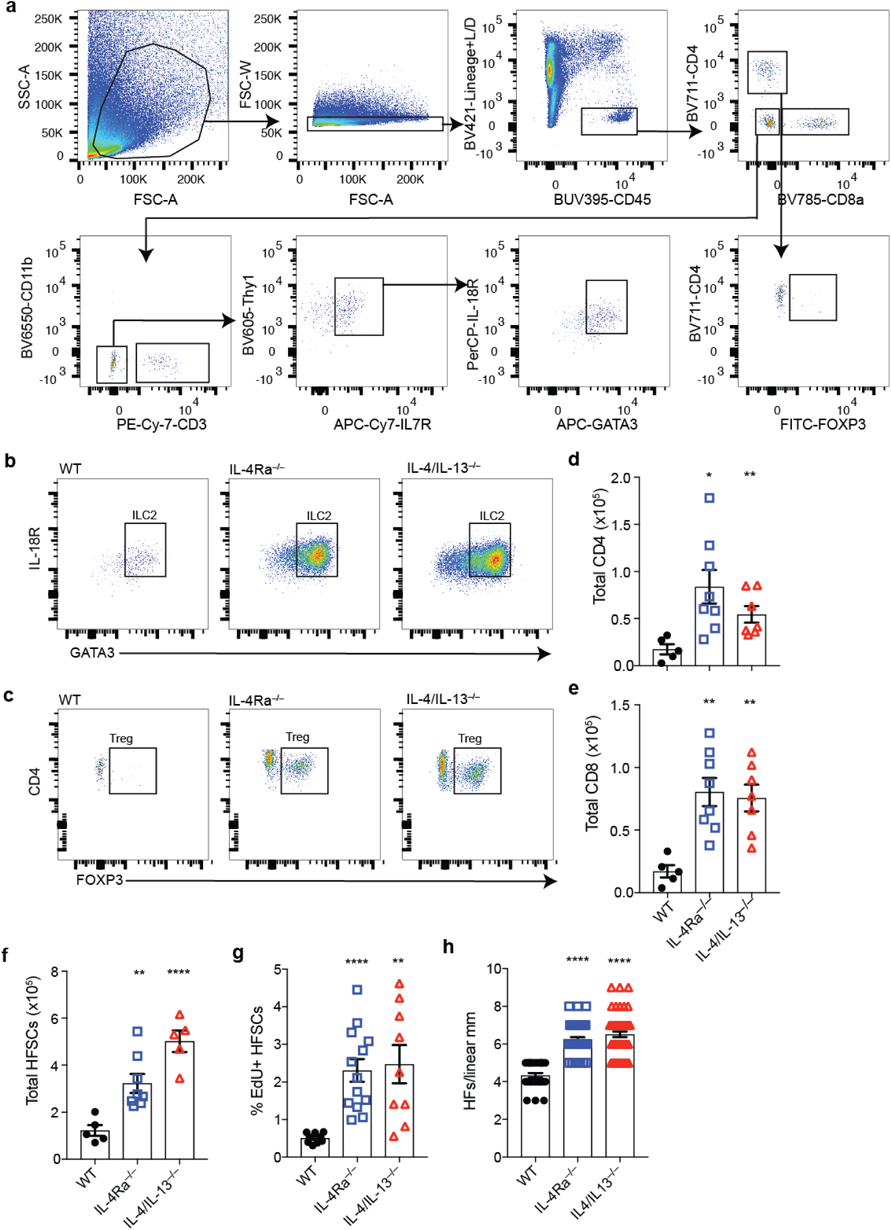
Type 2 immune-deficient strains have increased burden of *Demodex* mites. **a**, Gating strategy for skin cell populations using various mouse strains. **b**, Flow cytometry of skin ILC2s (CD45+Lin–Thy1+IL-7R+GATA3+) in 8-12 weeks old WT, IL-4Ra^−/−^, and IL-4/IL-13^−/−^ mice at homeostasis. **c**, Flow cytometry of skin regulatory T cells (Tregs) (CD3+CD4+FoxP3+) in 8-12 weeks old WT, IL-4Ra^−/−^, and IL-4/IL-13^−/−^ mice at homeostasis. **d**, Total CD4 in skin from WT, IL-4Ra^−/−^, and IL-4/IL-13^−/−^ mice. **e**, Total CD8 in skin from WT, IL-4Ra^−/−^, and IL-4/IL-13^−/−^ mice. **f**, Number of hair follicle stem cells. **g**, Percent EdU^+^ hair follicle stem cells (HFSCs). **h**, Quantification of visualized hair follicles per 1mm of skin. Data presented as mean ± s.e.m. Data are from one representative experiment (**a, d, e**) of at least two independent experiments, or pooled from multiple independent experiments (**b, c**). * P < 0.05, ** P < 0.01, *** P < 0.001, **** P < 0.0001 by two-tailed Student’s t test.

**Supplementary Fig. 4.**
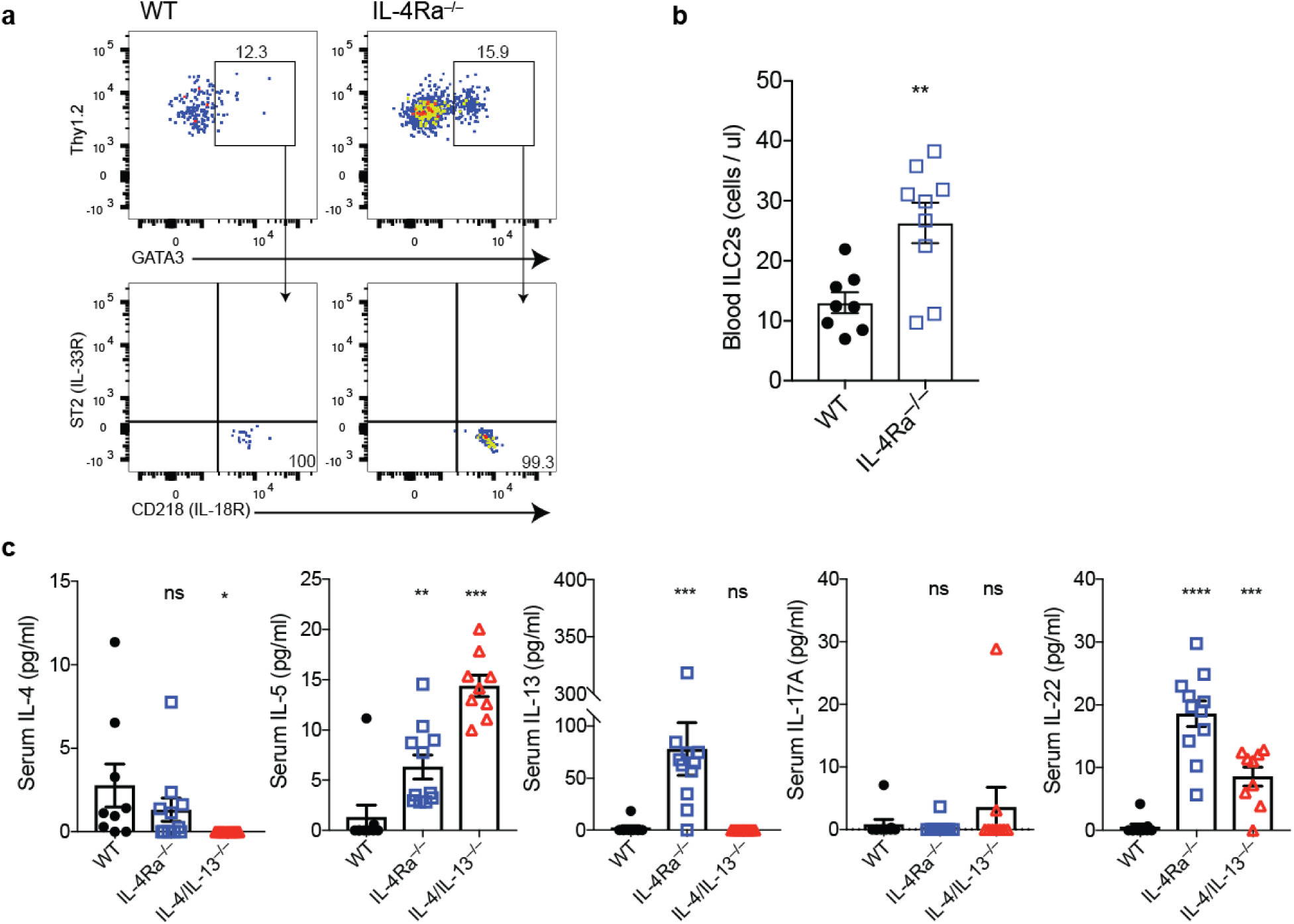
Activation and expansion of skin ILC2s leads to extrusion into the circulation. **a**, Flow cytometry plot for blood ILC2s (gated CD45^+^Lin^−^Thy1^+^GATA3^+^) in WT or *Demodex* infested IL-4Ra^−/−^ mice. Expression of ST2 and IL-18R in gated ILC2s as indicated. **b**, Frequency of blood ILC2s in WT or Demodex infested IL-4Ra^−/−^ mice. **c**, Serum IL-4, IL-5, IL-13, IL-17A, IL-22 in WT and Demodex infested IL-4Ra^−/−^ and IL-4/IL-13^−/−^ mice. Data presented as mean ± s.e.m and representative (**a**) or pooled (**b, c**) from at least two independent experiments. Statistical significance shown by * P < 0.05, ** P < 0.01, *** P < 0.001, **** P < 0.0001 by two-tailed Student’s t test.

**Supplementary Fig. 5.**
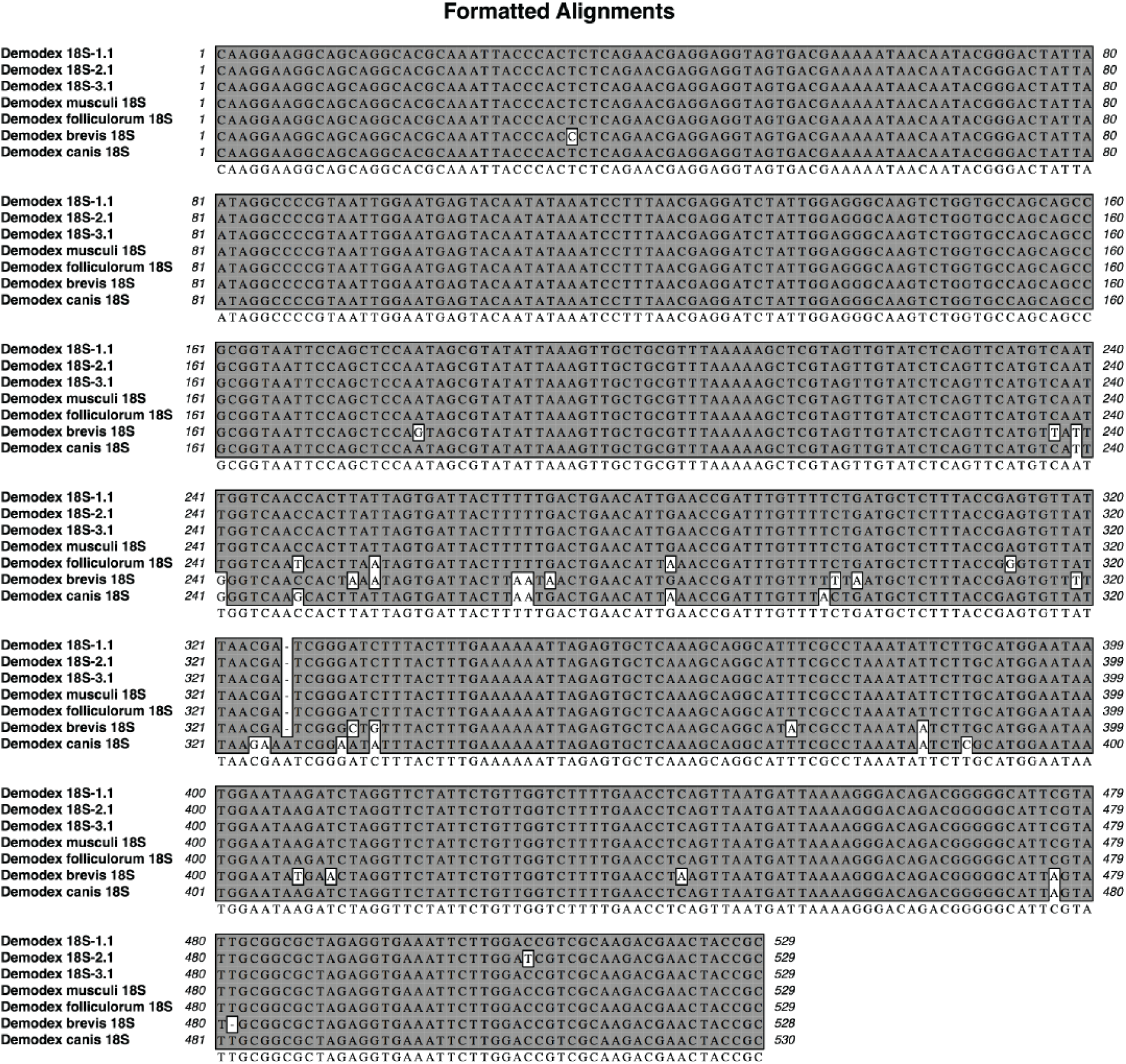
Identification of Demodex musculi. Alignments of putative *Demodex spp.* 18S rRNA gene amplicon sequences obtained from 3 affected mice (one representative for each sample) with the corresponding sequences of *Demodex musculi* (GenBank #JF834894), *Demodex folliculorum* (GenBank #KY922187), *Demodex brevis* (GenBank JN885466.1), and *Demodex canis* (GenBank JN885468.1). Single nucleotide sequence which differs from the *Demodex musculi* reference is highlighted and the consensus sequence is also shown.

**Supplementary Fig. 6.**
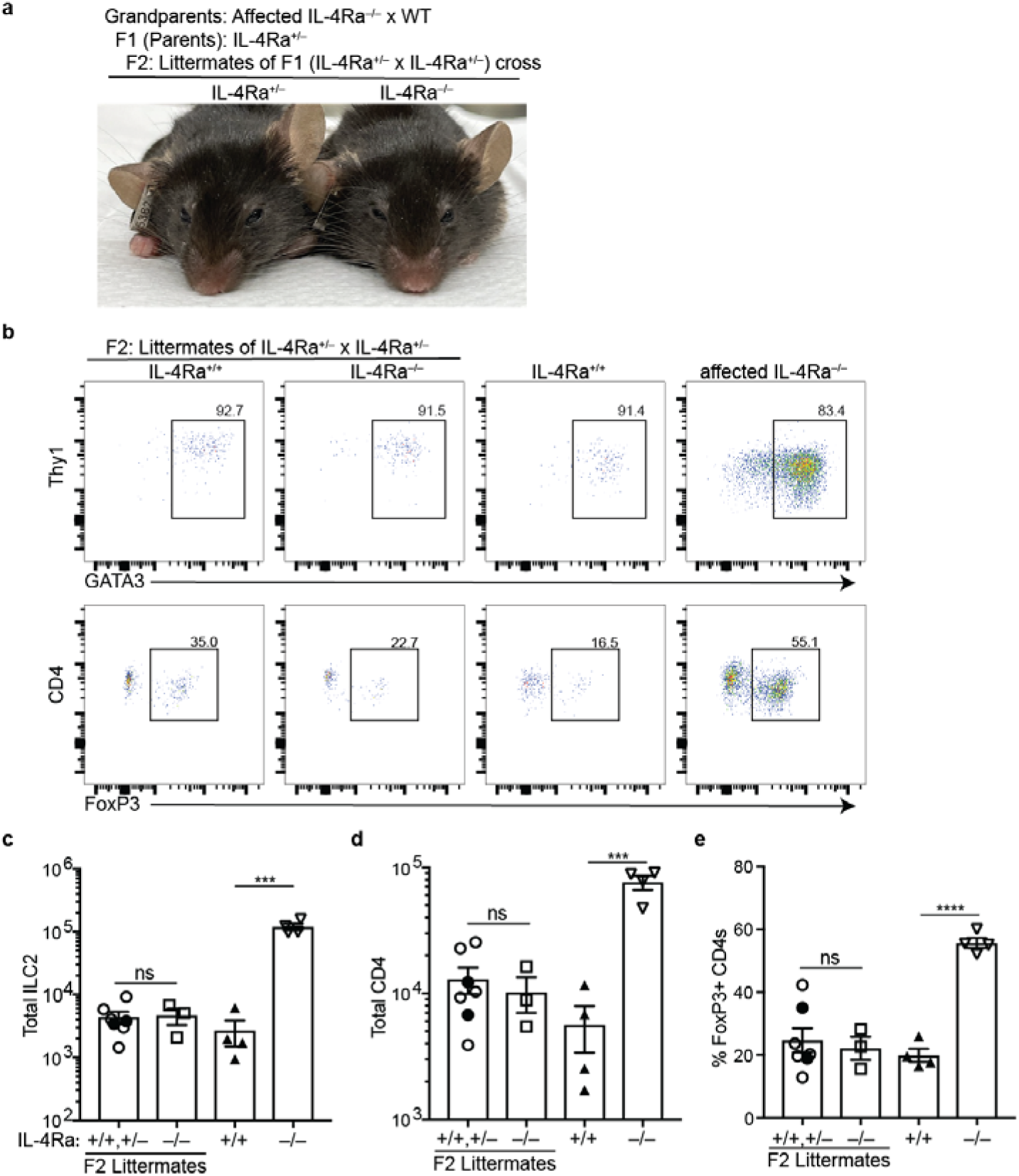
IL-4Ra^−/−^ mice lose the skin phenotype when derived through a generation with sufficient type 2 immunity. **a**, Facial picture of IL-4Ra^+/+^ and IL-4Ra^−/−^ F2 mice generated from F1 intercross of IL-4Ra^+/–^ parent mice. The F1 IL-4Ra^+/–^ mice were derived by crossing unaffected WT x affected IL-4Ra^−/−^ grandparents. **b**, Flow cytometry plots of ILC2s (top, gated CD45+Lin–CD3–CD4– Thy1+IL7R+) or CD4 (bottom, gated CD45+CD3+CD4+) from F2 IL-4Ra^+/+^, and IL-4Ra^−/−^ mice generated from F1 IL-4Ra^+/–^ intercross or age matched WT and affected IL-4Ra^−/−^ experimental groups. **c**–**e**, Quantification of ILC2 (**c**), CD4 (**d**), and frequency of Tregs (**e**). In the IL-4Ra^+/+^, and IL-4Ra^+/–^ experimental group, closed circles represent WT mice and open circles represent heterozygous mice. Data presented as mean ± s.e.m and representative of two independent experiments. Statistical significance shown by * P < 0.05, ** P < 0.01, *** P < 0.001, **** P < 0.0001 by one-way ANOVA.

**Supplementary Fig. 7.**
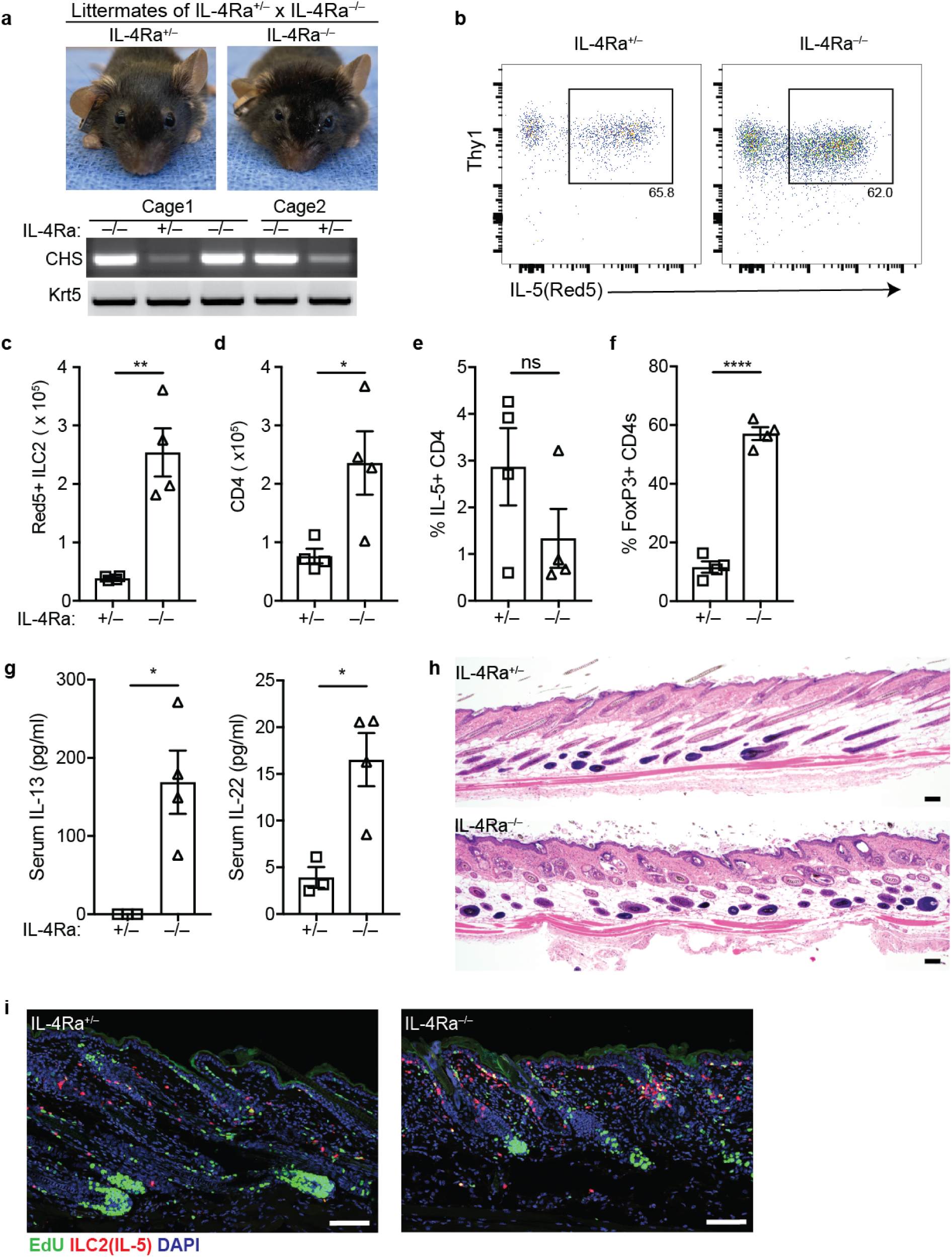
Mice with intact type 2 immunity do not show the altered skin phenotype associated with *Demodex* infection. **a**, Facial pictures of IL-4Ra^+/–^;IL-5^Red5^ and IL-4Ra^−/−^;IL-5^Red5^ littermates generated from visually unaffected IL-4Ra^+/–^ x affected IL-4Ra^−/−^ mice. The F1 IL-4Ra^+/–^ mice were derived by crossing unaffected WT x affected IL-4Ra^−/−^ parents. Bottom, PCR for *Demodex* chitinase synthase gene (CHS) or genomic DNA for the keratin 5 gene (Krt5) from back skin. **b**, Flow cytometry plots of Red5^+^ ILC2s (pre-gated on Live CD45^+^Lin^−^Thy1^+^) from littermate IL-4Ra^+/–^ and IL-4Ra^−/−^ mice. **c,d**, Quantification of Red5+ ILC2s (C) and CD4 (D) from back skin. **e**, Frequency of IL-5+(Red5+) CD4 T cells in skin. **f**, Frequency Tregs (CD3+CD4+FoxP3+) in skin. **g**, Serum IL-13 and IL-22 from littermate IL-4Ra^+/–^ and IL-4Ra^−/−^ mice. **h**, Skin sections stained with H&E. Scale bar, 100□m. **i**, Back skin sections were stained with EdU (green), anti-tdTomato (red, highlighting IL-5+ cells), and DAPI blue). Scale bar, 100 μm. Data presented as mean ± s.e.m and representative of two independent experiments. Statistical significance shown by * P < 0.05, ** P < 0.01, *** P < 0.001, **** P < 0.0001 by two-tailed Student’s t test. ns, not significant.

**Supplementary Fig. 8.**
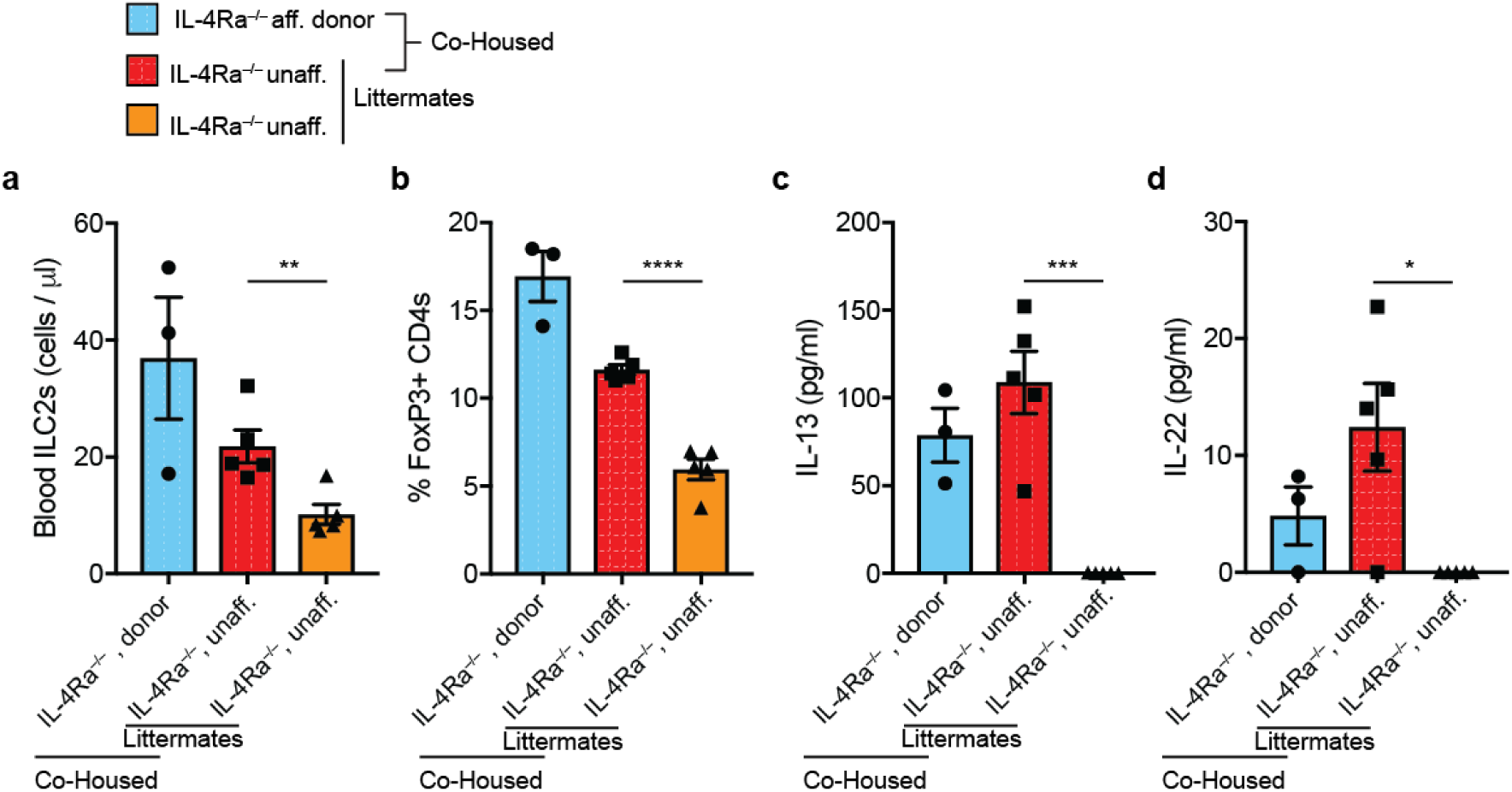
Co-housing *Demodex* infested and non-infested IL-4Ra^−/−^ mice transfers the blood phenotype. **a,b**, Frequency of circulating ILC2s (**a**) and Tregs (**b**) in IL-4Ra^−/−^ mice with or without Demodex (See Figure 4B). **c**,**d**, Serum IL-13 (**c**) and IL-22 (**d**). Data presented as mean ± s.e.m and representative of two independent experiments. Statistical significance shown by * P < 0.05, ** P < 0.01, *** P < 0.001, **** P < 0.0001 by two-tailed Student’s t test.

**Supplementary Fig. 9.**
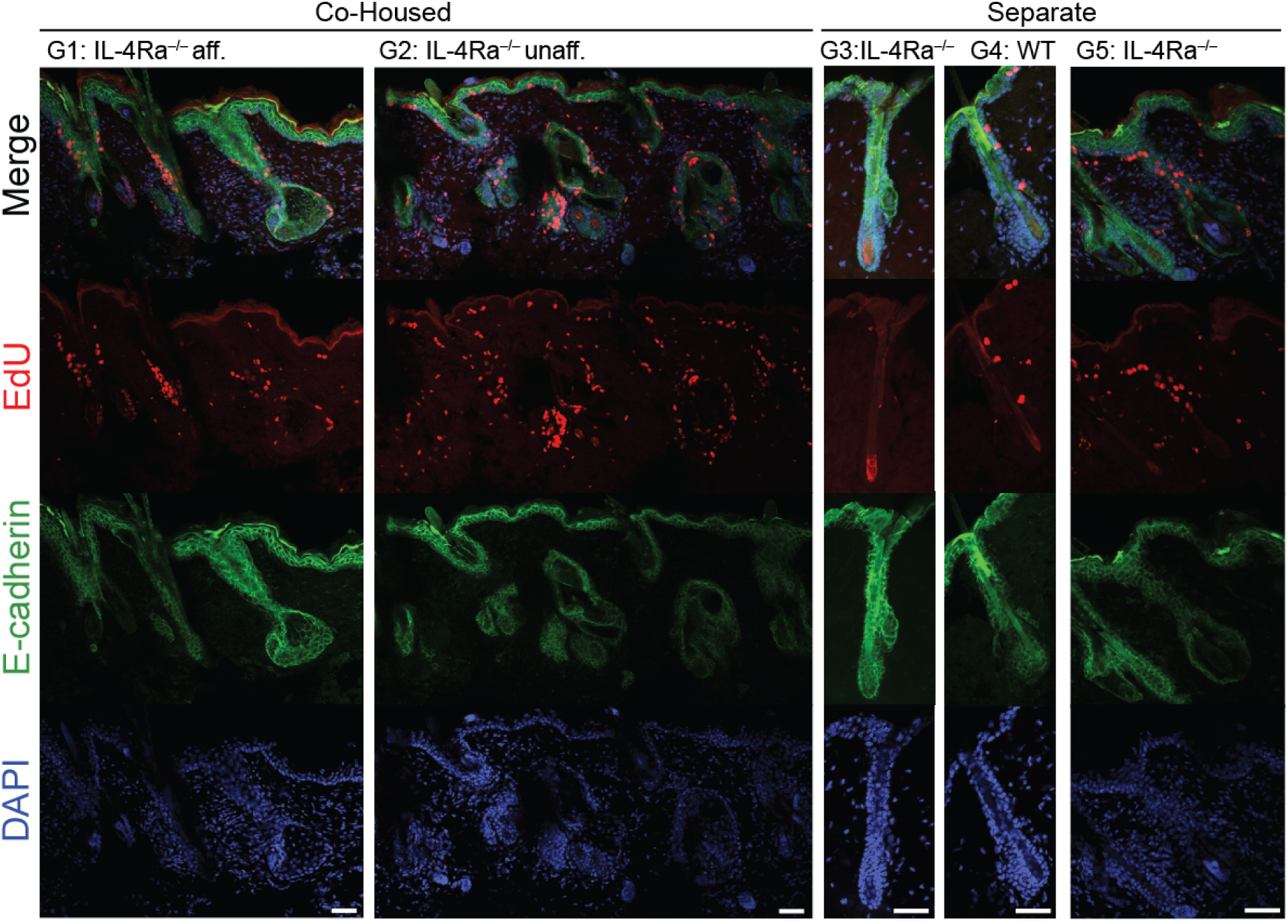
Epithelial proliferation in affected IL-4Ra^−/−^ mice. Representative sections from back skin of the co-housing experiment shown in Figure 4. 8 weeks old (w.o.) IL-4Ra^−/−^ mice infected with *Demodex* (Group 1) were co-housed with unaffected 3 w.o. IL-4Ra^−/−^ mice (Group 2). Unaffected littermates of co-housed mice (Group 3) were allowed to age concurrently. WT (Group 4) and IL-4Ra^−/−^ from known Demodex infected parents (Group 5) were aged as independent controls. Mice were injected with EdU 16 hours before harvest, and skin section were stained for EdU (Red), E-cadherin (green), and DAPI (blue). Scale bar, 100 μm.

**Supplementary Fig. 10.**
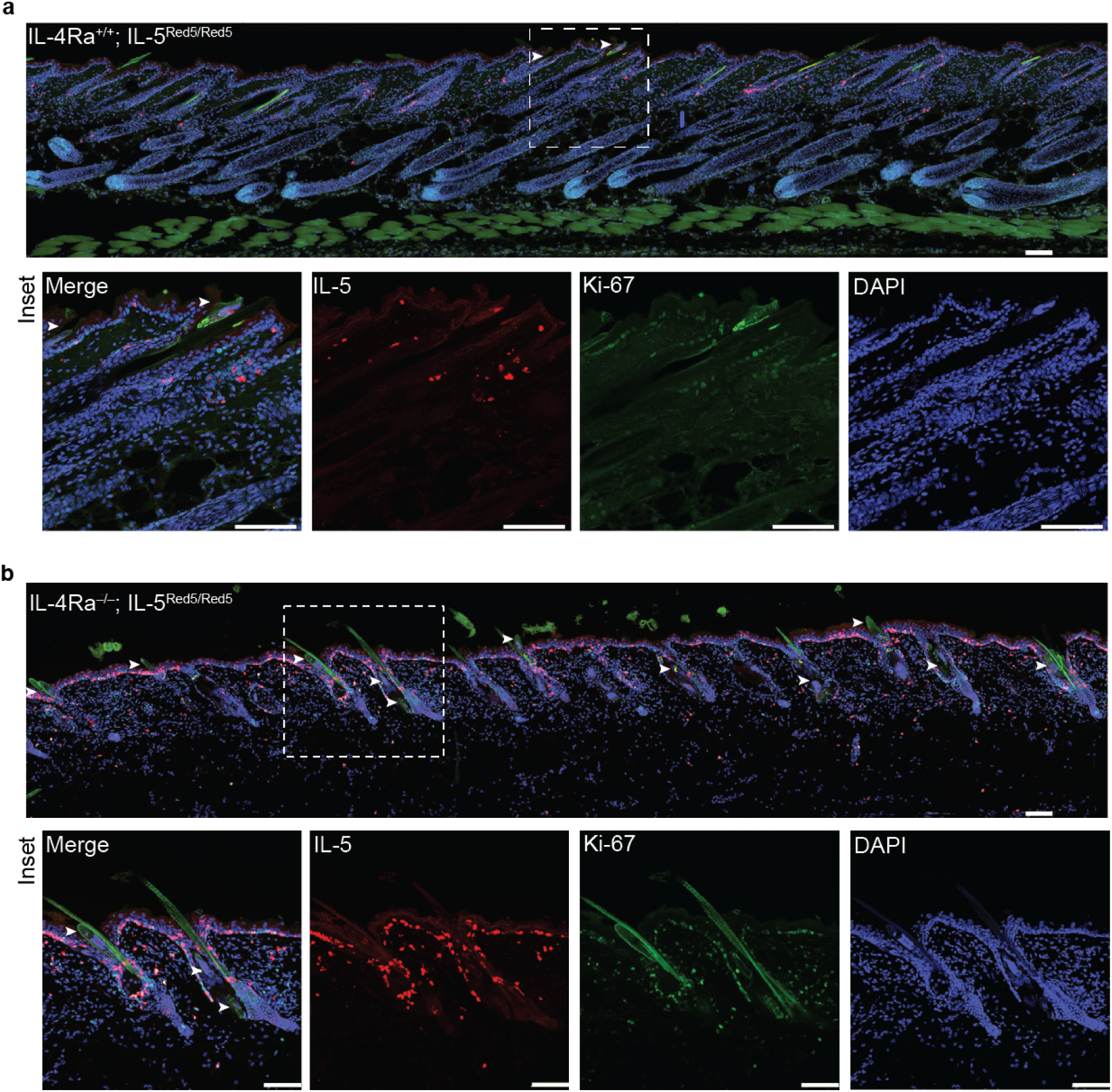
Increased number of IL-5-expressing ILC2s in IL-4Ra^−/−^ mice infected with *Demodex*. **a,b**, Representative images from back skin sections of IL-4Ra^+/+^;IL-5^Red5^ mice (**a**) co-housed with IL-4Ra^−/−^;IL-5^Red5^ mice (**b**) for 8 weeks were stained for tdTomato (IL-5 producing cells, red), Ki-67 (green), and DAPI (blue). The arrowheads highlight visible *Demodex* in the hair follicle or sebaceous glands. Scale bar, 100 μm.

**Supplementary Fig. 11.**
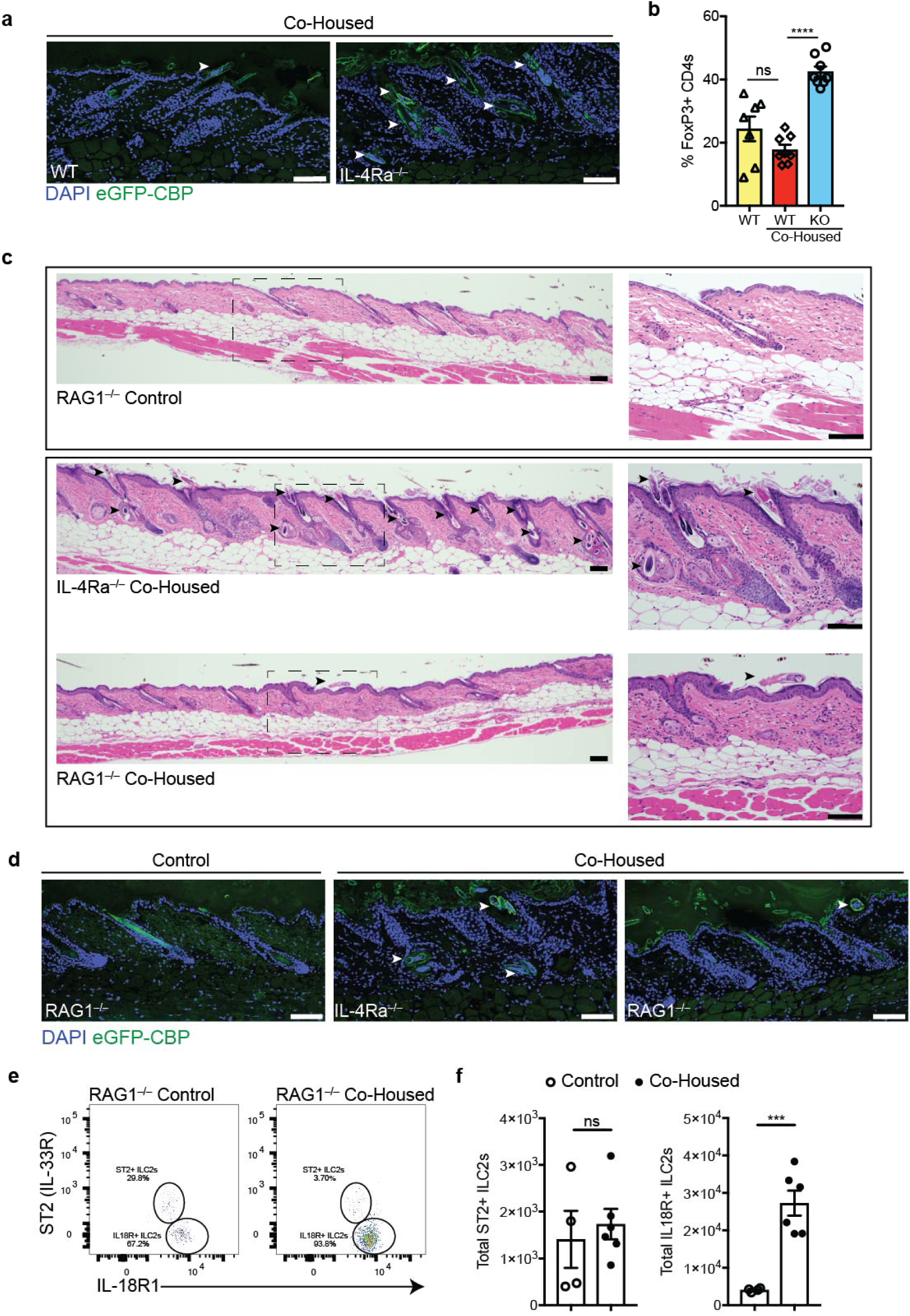
Type 2 innate immunity restrains *Demodex* infection. **a**, Skin sections from co-housed WT or *Demodex* infected IL-4Ra^−/−^ were stained for chitin (eGFP-CBP, green) and DAPI (blue). **b**, Frequency of Tregs (CD3+CD4+FoxP3+ T cells) as a percentage of total CD4 from back skin. **c**, Skin sections from RAG1^−/−^;IL-5^Red5/Red5^ control mice and demodex infected IL-4Ra^−/−^ mice co-housed with RAG1^−/−^;IL-5^Red5/Red5^ mice were stained with H&E. **d,** Skin sections as in (**c**), stained for chitin (green, eGFP-CBP) and DAPI (blue). In (**a**), (**c**) and (**d**), arrowheads highlight Demodex mites. All scale bars, 100 μm. **e**, Flow cytometry plots showing ST2 and IL-18R expression by skin ILC2s (pre-gated on Live CD45^+^Lin^−^Thy1^+^Red5^+^) from RAG1^−/−^;IL-5^Red5/Red5^ (control) or RAG1^−/−^;IL-5^Red5/Red5^ that were co-housed with IL-4Ra^−/−^ mice (Co-Housed). **f**, Quantification of the total of ST2+ (left) or IL-18R+ (right) Red5+ ILC2s in full thickness back skin. Data presented as mean ± s.e.m and pooled from 2 independent cohorts. Statistical significance shown by *** P < 0.001 by two-tailed Student’s t test. ns, not significant.

**Supplementary Fig. 12.**
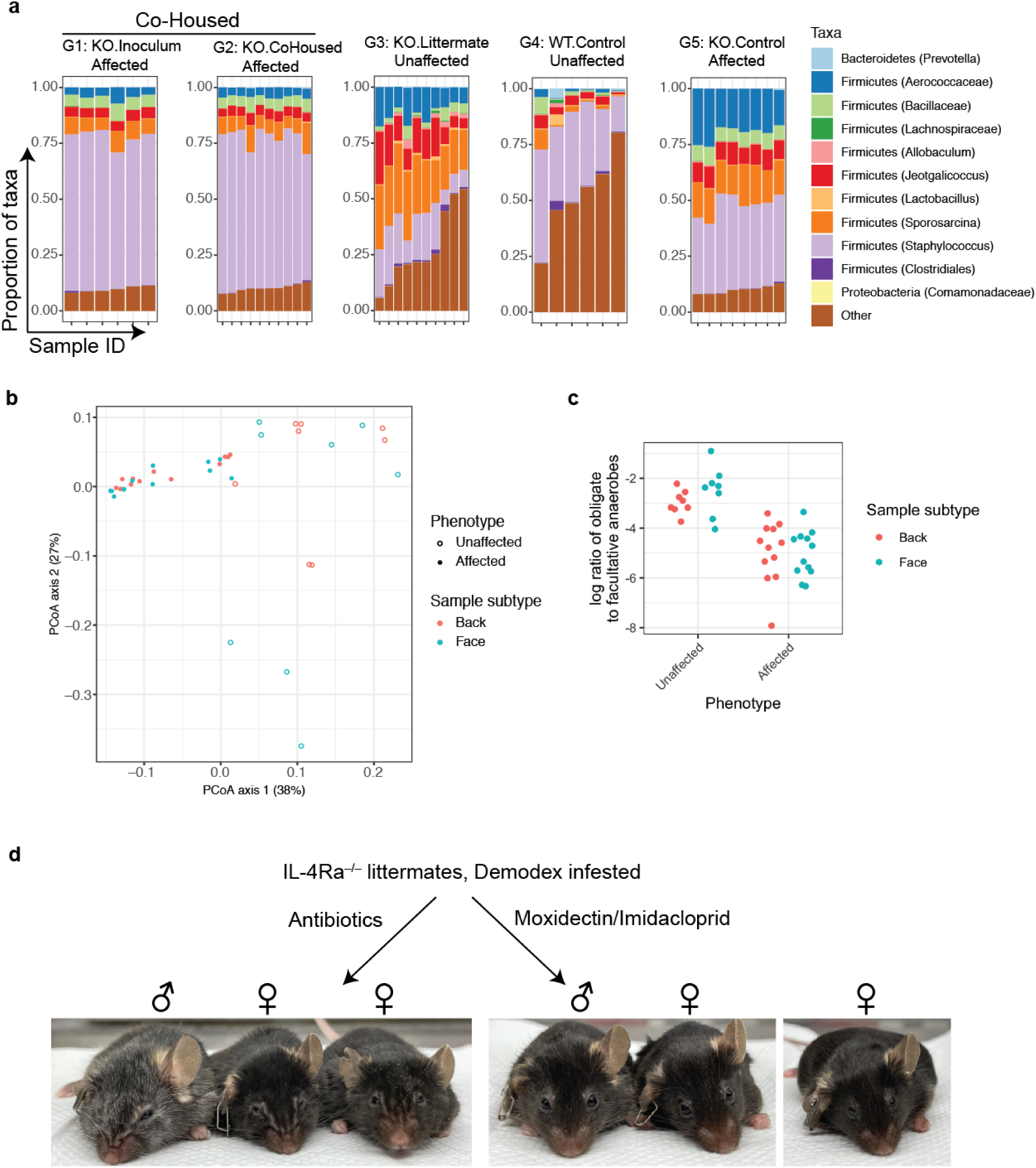
16S ribosomal RNA gene sequencing analysis of mice showing affected and unaffected skin phenotypes. **a**, Compositional plots of the top 10 major taxa present in each sample of the co-housing experiment shown in Figure 4. Bacterial lineages are color coded and presented as stacked bar graphs. **b**, PCoA plot based on the Bray-Curtis dissimilarity assessed using the microbiome data. All samples were used to make a common plot. Samples were further classified by gross skin phenotype (unaffected, open circles; affected, closed circles). The color of the symbol indicates the sampling site. **c**, Ratio of obligate to facultative anaerobes abundance in unaffected and affected mice. **d**, 8 week old, *Demodex* infested IL-4Ra^−/−^ littermates were separated and treated with broad spectrum antibiotics for 1 month in the drinking water or topically with Moxidectin/Imidacloprid weekly for 8 weeks.

**Supplementary Fig. 13.**
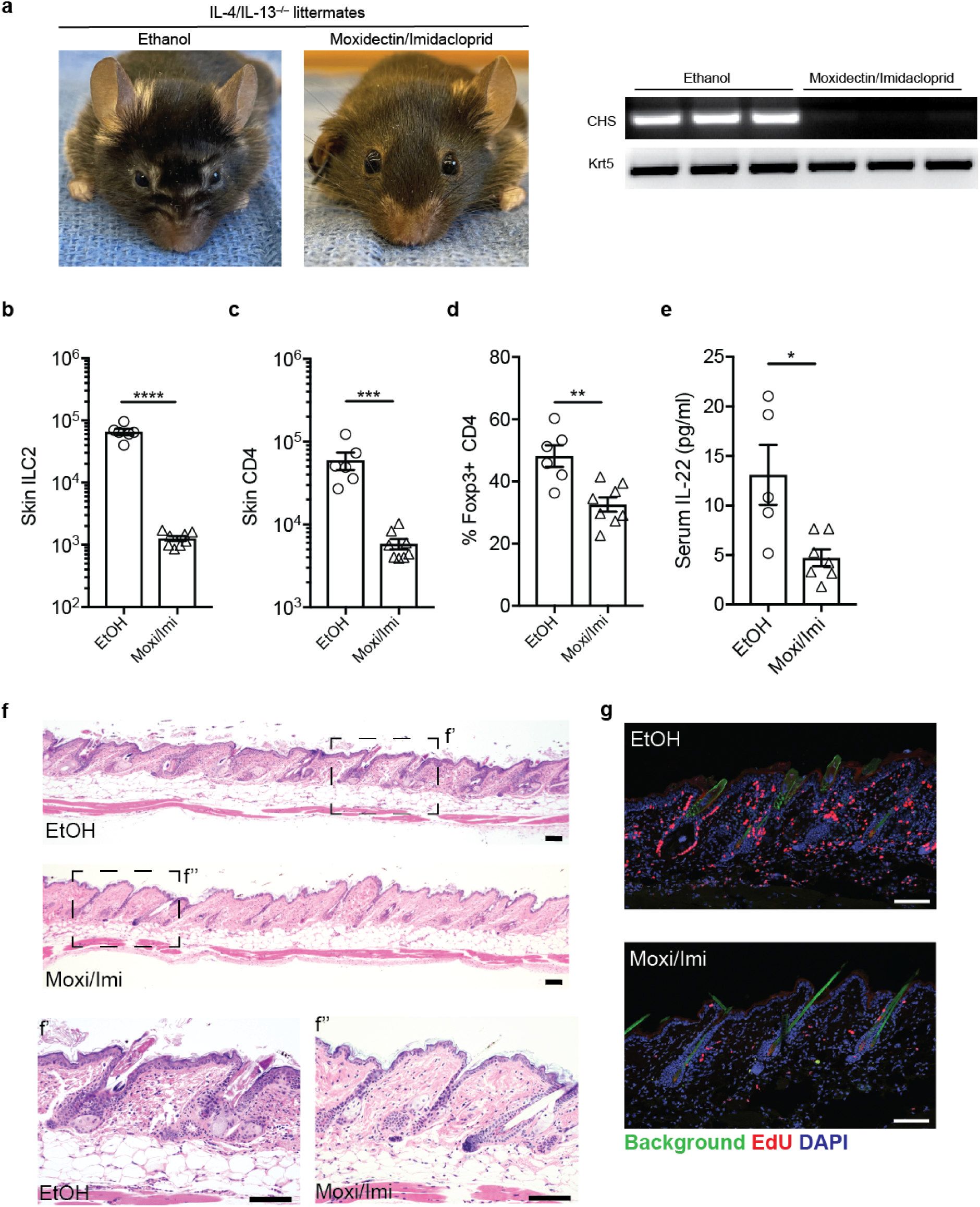
Targeted therapy reverses the phenotype in *Demodex* infested IL-4/IL-13^−/−^ mice. **a**, Facial pictures of IL-4/IL-13^−/−^ mice. Littermates of known *Demodex* infested IL-4/IL-13^−/−^ were separated at 3-6 weeks of age and treated with topical Ethanol (vehicle) or anti-parasitic Moxidectin/Imidacloprid (Moxi/Imi) once a week for 8 weeks. Right, representative PCR for *Demodex* chitinase synthase gene (CHS) or genomic DNA for the keratin 5 gene (Krt5) from back skin. **b**,**c**, Quantification of ILC2s (B) and CD4s (C) in IL-4/IL-13^−/−^ mice treated with Ethanol (EtOH) or Moxidectin/Imidacloprid (Moxi/Imi) for 8 weeks. **d**, Frequency of Tregs in IL-4/IL-13^−/−^ mice treated with EtOH or Moxi/Imi. **e**, Serum IL-22. **F**, Sections from back skin of IL-4/IL-13^−/−^ control (EtOH) or moxidectin/imidacloprid (Moxi/Imi) treated mice were stained with H&E. Scale bar, 100 μm. **g**, Sections from back skin of IL-4/IL-13^−/−^ control (EtOH) or moxidectin/imidacloprid (Moxi/Imi) were stained for EdU (red) and DAPI (blue). Scale bar, 100 μm. Data presented as mean ± s.e.m and pooled from two independent experiments. Statistical significance shown by ** P < 0.01, *** P < 0.001, **** P < 0.0001 by two-tailed Student’s t test.

**Supplementary Table 1.** List of antibodies used for flow cytometry and immunofluorescence.

| Company | Catalog Number | Antibody Name | Clone | Dilution |
| --- | --- | --- | --- | --- |
| BioLegend | 115549 | Brilliant Violet 421™ anti-mouse CD19 Antibody, 50 µg | 6D5 | 1:300 |
| BioLegend | 116234 | Brilliant Violet 421™ anti-mouse TER-119/Erythroid Cells Antibody, 50 µg | TER-119 | 1:300 |
| Biolegend | 108445 | Brilliant Violet 421™ anti-mouse Ly-6G/Ly-6C (Gr-1) Antibody, 50 µg | RB6-8C5 | 1:300 |
| Biolegend | 101236 | Brilliant Violet 421™ anti-mouse/human CD11b Antibody, 500ul | M1/70 | 1:300 |
| Biolegend | 101259 | Brilliant Violet 650™ anti-mouse/human CD11b Antibody | M1/70 | 1:300 |
| Biolegend | 117322 | Pacific Blue™ anti-mouse CD11c Antibody, 100 µg | N418 | 1:300 |
| BioLegend | 108918 | Pacific Blue™ anti-mouse CD49b (pan-NK cells) Antibody, 100 µg | DX5 | 1:300 |
| BioLegend | 137612 | Brilliant Violet 421™ anti-mouse CD335 (NKp46) Antibody, 50 µg | 29A1.4 | 1:300 |
| BioLegend | 108741 | Brilliant Violet 421™ anti-mouse NK-1.1 Antibody, 50 µg | PK136 | 1:300 |
| BioLegend | 118120 | Brilliant Violet 421™ anti-mouse TCR γ/δ Antibody, 50 µg | GL3 | 1:500 |
| Biolegend | 134314 | Pacific Blue™ anti-mouse FcεRIα Antibody, 100 µg | MAR-1 | 1:300 |
| BioLegend | 100220 | PE/Cy7 anti-mouse CD3 Antibody, 100 µg | 17A2 | 1:200 |
| BioLegend | 123124 | Pacific Blue™ anti-mouse F4/80 Antibody, 100 µg | BM8 | 1:300 |
| Biolegend | 103147 | Brilliant Violet 711™ anti-mouse CD45 Antibody, 50 µg | 30-F11 | 1:300 |
| BD | 564279 | BUV395 Rat Anti-Mouse CD45, 50 µg | 30-F11 | 1:300 |
| BioLegend | 103116 | APC/Cyanine7 anti-mouse CD45 Antibody, 100 µg | 30-F11 | 1:400 |
| BioLegend | 105331 | Brilliant Violet 785™ anti-mouse CD90.2 Antibody, 50 µg | 30-H12 | 1:1000 |
| Biolegend | 105343 | Brilliant Violet 605™ anti-mouse CD90.2 Antibody | 30-H12 | 1:1000 |
| BioLegend | 100550 | Brilliant Violet 711™ anti-mouse CD4 Antibody, 500 µl | RM4-5 | 1:200 |
| BioLegend | 100725 | Pacific Blue™ anti-mouse CD8a Antibody, 100 µg | 53-6.7 | 1:200 |
| BioLegend | 100750 | Brilliant Violet 785™ anti-mouse CD8a Antibody | 53-6.7 | 1:200 |
| BioLegend | 100642 | Pacific Blue™ anti-mouse CD5 Antibody | 53-7.3 | 1:300 |
| BioLegend | 108133 | Brilliant Violet 605™ anti-mouse Ly-6A/E (Sca-1) Antibody, 125 µL | D7 | 1:300 |
| BioLegend | 107643 | Brilliant Violet 711™ anti-mouse I-A/I-E Antibody | M5/114.15.2 | 1:400 |
| BioLegend | 118220 | PerCP/Cyanine5.5 anti-mouse CD326 (Ep-CAM) Antibody | G8.8 | 1:300 |
| BioLegend | 313606 | FITC anti-human/mouse CD49f Antibody | GoH3 | 1:300 |
| BioLegend | 313622 | PE/Cyanine7 anti-human/mouse CD49f Antibody | GoH3 | 1:300 |
| BD | 560230 | Alexa Fluor® 647 Rat anti-Mouse CD34 | RAM34 | 1:200 |
| Invitrogen | 47-1271-82 | CD127 (IL-7R) Monoclonal Antibody (A7R34), APC-eFluor 780 | A7R34 | 1:100 |
| Invitrogen | 50-9966-42 | Gata-3 Monoclonal Antibody (TWAJ), eFluor 660, eBioscience™, 100 tests | TWAJ | 1:25 |
| Biolegend | 300514 | APC anti-human CD4 Antibody | RPA-T4 | 1:20 |
| Invitrogen | 11-5773-82 | FOXP3 Monoclonal Antibody (FJK-16s), eBioscience™ | FJK-16s | 1:100 |
| BD | 745257 | BV605 Rat Anti-Mouse IL-33R (ST2) , 50 µg | U29-93 | 1:200 |
| Invitrogen | 46-5183-82 | CD218a (IL-18Ra) Monoclonal Antibody (P3TUNYA), PerCP-eFluor 710, eBioscience™, 100 µg | P3TUNYA | 1:300 |
| TakaraBio | 632496 | Living Colors® DsRed Polyclonal Antibody (100 µl) | Polyclonal, Lot 1904182 | 1:500 |
| Invitrogen | 11-5698-82 | Ki-67 Monoclonal Antibody (SolA15), FITC, eBioscience™ | SolA15 | 1:100 |
| BD | 612131 | FITC Mouse Anti- E-Cadherin | Clone 36 | 1:400 |
| Invitrogen | A-21428 | Goat anti-Rabbit IgG (H+L) Cross-Adsorbed Secondary Antibody, Alexa Fluor 555 | Polyclonal, Lot 2011559 | 1:1000 |

**Supplementary Table 2.** List of antibodies used for the CyTOF experiment.

| Channel | Metal | Antigen | Conc. Stock<br>(mg/ml) | Clone | Vendor |
| --- | --- | --- | --- | --- | --- |
| 113 | In | Ter119 | 0.4 | TER119 | Biolegend |
| 115 | In | CD45 | 0.5 | 30-F11 | Biolegend |
| 139 | La | pSTAT3 | 3 | 4 | BD |
| 140 | Ce | CD326 | 0.5 | G8.8 | Biolegend |
| 141 | Pr | CD11b | 0.5 | M1/70 | Biolegend |
| 142 | Nd | pH-H3 | 0.5 | HTA28 | Abcam |
| 143 | Nd | CD11c | 0.2 | HL3 | BD |
| 144 | Nd | pMAPKAPK2 | 1.5 | 27B7 | CST |
| 145 | Nd | pCREB | 4 | 87G3 | CST |
| 146 | Nd | pPLCg2 | 1 | K86-689.37 | BD |
| 147 | Sm | pSTAT1 | 2.5 | 4a | BD |
| 148 | Nd | Myc | 0.2 | 9E10 | Fluidigm |
| 149 | Sm | pSTAT5 | 3 | 47 | BD |
| 150 | Nd | pS6 | 7 | 2F9 | CST |
| 151 | Eu | Ly6C | 0.5 | HK1.4 | Biolegend |
| 152 | Sm | pErk1/2 | 10 | D13.14.4E | CST |
| 153 | Eu | CyclinB1-153 | 0.2 | GNS-1 | Fluidigm |
| 154 | Sm | p-p38 | 3 | 36/p38 | BD |
| 155 | Gd | CD8 | 0.2 | 53-6.7 | Biolegend |
| 156 | Gd | CD4 | 0.5 | RM4-5 | Biolegend |
| 157 | Gd | CD3 | 0.5 | 17A2 | BD |
| 158 | Gd | Ly6G | 0.5 | 1A8-FDA13 | Biolegend |
| 159 | Tb | p4EBP1 | 1 | 136B4 | CST |
| 160 | Gd | pSyk/ZAP70 | 1.25 | 17a | BD |
| 161 | Dy | pTBK1 | 6 | D52C2 | CST |
| 162 | Dy | TCRgd | 0.853 | GL3 | Biolegend |
| 163 | Dy | IkBα | 3 | L35A5 | CST |
| 164 | Dy | AFP-1 | 0.2 | SPM334 | Novus |
| 165 | Ho | pSTAT6 | 3 | 18 | DB |
| 166 | Er | pRB | 0.2 | J112906 | Fluidigm |
| 167 | Er | Foxp3 | 5 | NRRF-30 | ebioscience |
| 168 | Er | NK1.1 | 0.5 | PK136 | Biolegend |
| 169 | Tm | Ki67 | 2 | SolA15 | eBioscience |
| 170 | Er | pSTAT4 | 3 | 38 | BD |
| 171 | Yb | CD62L fitc | 5 | FIT-22 | Biolegend |
| 172 | Yb | SiglecF | 0.5 | E50-2440 | BD |
| 173 | Yb | CD19 | 0.5 | 6D5 | Biolegend |
| 174 | Yb | CD34 | 0.2 | RAM34 | BD |
| 175 | Lu | CD44 | 0.5 | IM7 | BD |
| 209 | Bi | MHC II | 0.5 | M5/114.15.2 | Biolegend |

## References

1 Hammad, H. & Lambrecht, B. N. Barrier Epithelial Cells and the Control of Type 2 Immunity. Immunity 43, 29–40, doi:10.1016/j.immuni.2015.07.007 (2015).

2 Gieseck, R. L., 3rd, Wilson, M. S. & Wynn, T. A. Type 2 immunity in tissue repair and fibrosis. Nat Rev Immunol 18, 62-76, doi:10.1038/nri.2017.90 (2018).

3 Klose, C. S. N. & Artis, D. Innate lymphoid cells control signaling circuits to regulate tissue-specific immunity. Cell Res 30, 475–491, doi:10.1038/s41422-020-0323-8 (2020).

4 Guttman-Yassky, E. et al. Dupilumab progressively improves systemic and cutaneous abnormalities in patients with atopic dermatitis. J Allergy Clin Immunol 143, 155–172, doi:10.1016/j.jaci.2018.08.022 (2019).

5 Liang, H. E. et al. Divergent expression patterns of IL-4 and IL-13 define unique functions in allergic immunity. Nat Immunol 13, 58–66, doi:10.1038/ni.2182 (2011).

6 Nussbaum, J. C. et al. Type 2 innate lymphoid cells control eosinophil homeostasis. Nature 502, 245–248, doi:10.1038/nature12526 (2013).

7 Ricardo-Gonzalez, R. R. et al. Tissue signals imprint ILC2 identity with anticipatory function. Nat Immunol, doi:10.1038/s41590-018-0201-4 (2018).

8 Schneider, C. et al. Tissue-Resident Group 2 Innate Lymphoid Cells Differentiate by Layered Ontogeny and In Situ Perinatal Priming. Immunity, doi:10.1016/j.immuni.2019.04.019 (2019).

9 Naik, S., Larsen, S. B., Cowley, C. J. & Fuchs, E. Two to Tango: Dialog between Immunity and Stem Cells in Health and Disease. Cell 175, 908–920, doi:10.1016/j.cell.2018.08.071 (2018).

10 Ali, N. et al. Regulatory T Cells in Skin Facilitate Epithelial Stem Cell Differentiation. Cell 169, 1119–1129 e1111, doi:10.1016/j.cell.2017.05.002 (2017).

11 Paus, R. et al. A comprehensive guide for the recognition and classification of distinct stages of hair follicle morphogenesis. J Invest Dermatol 113, 523–532, doi:10.1046/j.1523-1747.1999.00740.x (1999).

12 Keyes, B. E. et al. Impaired Epidermal to Dendritic T Cell Signaling Slows Wound Repair in Aged Skin. Cell 167, 1323–1338 e1314, doi:10.1016/j.cell.2016.10.052 (2016).

13 Ricardo-Gonzalez, R. R. et al. Tissue-specific pathways extrude activated ILC2s to disseminate type 2 immunity. J Exp Med 217, doi:10.1084/jem.20191172 (2020).

14 Huang, Y. et al. S1P-dependent interorgan trafficking of group 2 innate lymphoid cells supports host defense. Science 359, 114–119, doi:10.1126/science.aam5809 (2018).

15 Campbell, L. et al. ILC2s mediate systemic innate protection by priming mucus production at distal mucosal sites. J Exp Med 216, 2714–2723, doi:10.1084/jem.20180610 (2019).

16 Reynolds, G. et al. Developmental cell programs are co-opted in inflammatory skin disease. Science 371, doi:10.1126/science.aba6500 (2021).

17 Bielecki, P. et al. Skin-resident innate lymphoid cells converge on a pathogenic effector state. Nature, doi:10.1038/s41586-021-03188-w (2021).

18 Sanman, L. E. et al. Transit-Amplifying Cells Coordinate Changes in Intestinal Epithelial Cell-Type Composition. Dev Cell, doi:10.1016/j.devcel.2020.12.020 (2021).

19 Rosshart, S. P. et al. Laboratory mice born to wild mice have natural microbiota and model human immune responses. Science 365, doi:10.1126/science.aaw4361 (2019).

20 Sastre, N. et al. Detection, Prevalence and Phylogenetic Relationships of Demodex spp and further Skin Prostigmata Mites (Acari, Arachnida) in Wild and Domestic Mammals. PLoS One 11, e0165765, doi:10.1371/journal.pone.0165765 (2016).

21 Zhao, Y. E. et al. Cloning and sequence analysis of chitin synthase gene fragments of Demodex mites. J Zhejiang Univ Sci B 13, 763–768, doi:10.1631/jzus.B1200155 (2012).

22 Kobayashi, T. et al. Homeostatic Control of Sebaceous Glands by Innate Lymphoid Cells Regulates Commensal Bacteria Equilibrium. Cell 176, 982–997 e916, doi:10.1016/j.cell.2018.12.031 (2019).

23 Nashat, M. A. et al. Ivermectin-compounded Feed Compared with Topical Moxidectin-Imidacloprid for Eradication of Demodex musculi in Laboratory Mice. J Am Assoc Lab Anim Sci 57, 483–497, doi:10.30802/AALAS-JAALAS-18-000003 (2018).

24 Smith, P. C., Zeiss, C. J., Beck, A. P. & Scholz, J. A. Demodex musculi Infestation in Genetically Immunomodulated Mice. Comp Med 66, 278–285 (2016).

25 Bernink, J. H. et al. c-Kit-positive ILC2s exhibit an ILC3-like signature that may contribute to IL-17-mediated pathologies. Nat Immunol 20, 992–1003, doi:10.1038/s41590-019-0423-0 (2019).

26 Harrison, O. J. et al. Commensal-specific T cell plasticity promotes rapid tissue adaptation to injury. Science 363, doi:10.1126/science.aat6280 (2019).

27 Boniface, K. et al. IL-22 inhibits epidermal differentiation and induces proinflammatory gene expression and migration of human keratinocytes. J Immunol 174, 3695–3702, doi:10.4049/jimmunol.174.6.3695 (2005).

28 Schneider, C. et al. A Metabolite-Triggered Tuft Cell-ILC2 Circuit Drives Small Intestinal Remodeling. Cell 0, doi:10.1016/j.cell.2018.05.014 (2018).

29 Howitt, M. R. et al. Tuft cells, taste-chemosensory cells, orchestrate parasite type 2 immunity in the gut. Science 351, 1329–1333, doi:10.1126/science.aaf1648 (2016).

30 Palopoli, M. F. et al. Global divergence of the human follicle mite Demodex folliculorum: Persistent associations between host ancestry and mite lineages. Proc Natl Acad Sci U S A 112, 15958–15963, doi:10.1073/pnas.1512609112 (2015).

31 Thoemmes, M. S., Fergus, D. J., Urban, J., Trautwein, M. & Dunn, R. R. Ubiquity and diversity of human-associated Demodex mites. PLoS One 9, e106265, doi:10.1371/journal.pone.0106265 (2014).

32 Georgala, S. et al. Increased density of Demodex folliculorum and evidence of delayed hypersensitivity reaction in subjects with papulopustular rosacea. J Eur Acad Dermatol Venereol 15, 441–444, doi:10.1046/j.1468-3083.2001.00331.x (2001).

33 Zhao, Y. E., Wu, L. P., Peng, Y. & Cheng, H. Retrospective analysis of the association between Demodex infestation and rosacea. Arch Dermatol 146, 896–902, doi:10.1001/archdermatol.2010.196 (2010).

34 Rather, P. A. & Hassan, I. Human demodex mite: the versatile mite of dermatological importance. Indian J Dermatol 59, 60–66, doi:10.4103/0019-5154.123498 (2014).

35 Quint, T. et al. Dupilumab for the Treatment of Atopic Dermatitis in an Austrian Cohort-Real-Life Data Shows Rosacea-Like Folliculitis. J Clin Med 9, doi:10.3390/jcm9041241 (2020).

36 Bando, J. K., Nussbaum, J. C., Liang, H. E. & Locksley, R. M. Type 2 innate lymphoid cells constitutively express arginase-I in the naive and inflamed lung. J Leukoc Biol 94, 877–884, doi:10.1189/jlb.0213084 (2013).

37 von Moltke, J., Ji, M., Liang, H. E. & Locksley, R. M. Tuft-cell-derived IL-25 regulates an intestinal ILC2-epithelial response circuit. Nature 529, 221–225, doi:10.1038/nature16161 (2016).

38 Liang, H.-E. et al. Divergent expression patterns of IL-4 and IL-13 define unique functions in allergic immunity. Nature Immunology 13, 58–66, doi:10.1038/ni.2182 (2012).

39 Nagao, K. et al. Stress-induced production of chemokines by hair follicles regulates the trafficking of dendritic cells in skin. Nat Immunol 13, 744–752, doi:10.1038/ni.2353 (2012).

40 Kleppe, M. et al. Jak1 Integrates Cytokine Sensing to Regulate Hematopoietic Stem Cell Function and Stress Hematopoiesis. Cell Stem Cell 21, 489–501 e487, doi:10.1016/j.stem.2017.08.011 (2017).

41 Behbehani, G. K. et al. Transient partial permeabilization with saponin enables cellular barcoding prior to surface marker staining. Cytometry A 85, 1011–1019, doi:10.1002/cyto.a.22573 (2014).

42 Zunder, E. R., Lujan, E., Goltsev, Y., Wernig, M. & Nolan, G. P. A continuous molecular roadmap to iPSC reprogramming through progression analysis of single-cell mass cytometry. Cell Stem Cell 16, 323–337, doi:10.1016/j.stem.2015.01.015 (2015).

43 Finck, R. et al. Normalization of mass cytometry data with bead standards. Cytometry A 83, 483–494, doi:10.1002/cyto.a.22271 (2013).

44 Spitzer, M. H. et al. IMMUNOLOGY. An interactive reference framework for modeling a dynamic immune system. Science 349, 1259425, doi:10.1126/science.1259425 (2015).

45 Bair, E. & Tibshirani, R. Semi-supervised methods to predict patient survival from gene expression data. PLoS Biol 2, E108, doi:10.1371/journal.pbio.0020108 (2004).

46 Bruggner, R. V., Bodenmiller, B., Dill, D. L., Tibshirani, R. J. & Nolan, G. P. Automated identification of stratifying signatures in cellular subpopulations. Proc Natl Acad Sci U S A 111, E2770–2777, doi:10.1073/pnas.1408792111 (2014).

47 Spitzer, M. H. et al. Systemic Immunity Is Required for Effective Cancer Immunotherapy. Cell 168, 487–502 e415, doi:10.1016/j.cell.2016.12.022 (2017).

48 Bolyen, E. et al. Reproducible, interactive, scalable and extensible microbiome data science using QIIME 2. Nat Biotechnol 37, 852–857, doi:10.1038/s41587-019-0209-9 (2019).

49 Callahan, B. J. et al. DADA2: High-resolution sample inference from Illumina amplicon data. Nat Methods 13, 581–583, doi:10.1038/nmeth.3869 (2016).

50 McDonald, D. et al. An improved Greengenes taxonomy with explicit ranks for ecological and evolutionary analyses of bacteria and archaea. ISME J 6, 610–618, doi:10.1038/ismej.2011.139 (2012).

51 Bokulich, N. A. et al. Optimizing taxonomic classification of marker-gene amplicon sequences with QIIME 2’s q2-feature-classifier plugin. Microbiome 6, 90, doi:10.1186/s40168-018-0470-z (2018).

52 Katoh, K. & Standley, D. M. MAFFT multiple sequence alignment software version 7: improvements in performance and usability. Mol Biol Evol 30, 772–780, doi:10.1093/molbev/mst010 (2013).

53 Lozupone, C. & Knight, R. UniFrac: a new phylogenetic method for comparing microbial communities. Appl Environ Microbiol 71, 8228–8235, doi:10.1128/AEM.71.12.8228-8235.2005 (2005).

54 Lozupone, C. A., Hamady, M., Kelley, S. T. & Knight, R. Quantitative and qualitative beta diversity measures lead to different insights into factors that structure microbial communities. Appl Environ Microbiol 73, 1576–1585, doi:10.1128/AEM.01996-06 (2007).

55 Anderson, M. J. A new method for non-parametric multivariate analysis of variance. Austral Ecology 26, 32–46, doi:https://doi.org/10.1111/j.1442-9993.2001.01070.pp.x (2001).

